# Antibody-Protein L Functionalized Microparticles for Detection of Surface Markers in Heterogeneous Colorectal Lesions

**DOI:** 10.1101/2025.11.25.690572

**Authors:** Saleh Ramezani, Niki M. Zacharias, William Norton, Jennifer S. Davis, Abishai Dominic, Ryan Armijo, Muxin Wang, Richard E. Wendt, Daniel D. Carson, Daniel A. Harrington, Mary C. Farach-Carson, Pratip K. Bhattacharya

## Abstract

Visualization of colorectal cancer (CRC) lesions is complicated by their location in the colon and tumor morphology. Reliance on a single surface biomarker for direct identification risks false negatives due to temporal changes and/or tumor heterogeneity. We developed a multiplexed system of complementary biomarker targets in an effort to capture a broader range of lesions with diverse temporal and/or phenotypic expression. We identified Mucin-1 (MUC1) and epithelial cell adhesion molecule (EPCAM) as useful targeting pairs by examining multiple colon tumor subtypes in a standard tissue array, and by surveying multiple CRC cell lines, both as 2D cultures and as 3D tumoroids, for the presence of the CRC surface biomarkers. We demonstrated the utility of a “universal” surface functionalization approach using Antibody-Protein L functionalized microparticles (APL-MPs) that enabled the simultaneous incorporation of antibodies recognizing MUC1 and EPCAM. Using CRC cell heterogeneous tumoroids expressing both MUC1 and EPCAM (HET-tumoroids) and orthotopic animal cancer models designed to express both surface antigens, we demonstrated that: 1) APL-MPs identified MUC1- and EPCAM-positive tumoroids in proportion to antigen expression; 2) APL-MPs detected CRC surface antigens on the luminal colon surface *in vivo*, and 3) concurrent targeting of multiple surface antigens enhanced the sensitivity of detection of heterogeneous CRC lesions. This approach opens the door for the use of antibody–protein L dual-targeting MPs in a variety of applications to detect heterogeneous cancer lesions.

**Significance:** Multi-antigen targeting of MUC1 and EPCAM with Antibody-Protein L microparticles enhances the detection sensitivity of heterogeneous colorectal cancer lesions, offering a promising strategy for the accurate visualization of tumors in complex environments.

## 1. Introduction

Colorectal cancer (CRC) is a complex and heterogeneous disease with diverse phenotypic surface features. This heterogeneity poses a major challenge for the development of effective diagnostic tools and design of targeted therapies (1–4). Recent advances in imaging technology using high-affinity recognition molecules, such as antibodies, aptamers, peptides, or affibodies, have enhanced detection (5). Such molecules, when tethered to fluorescently labeled microparticles (MPs), can act as contrast agents that aid in visualization of malignant phenotypes (6–9); however, because of tumor heterogeneity, antibodies against single antigen targets do not possess sufficient sensitivity and specificity for reliable diagnostic purposes (10,11). To address current detection limitations, we developed a “universal” surface functionalization approach using Protein L-coated MPs that enables modular incorporation of multiple antibodies against different surface target antigens. Protein L is a bacterial protein that specifically binds to the kappa light chain of a wide range of immunoglobulin (Ig) classes, particularly IgG, IgM, IgD, IgA, Fab, and scF, without interfering with the antigen binding site (12). By leveraging the extensive binding capabilities of Protein L, we created a versatile system that combines fluorophore-labeled antibodies with Protein L to target and recognize multiple antigens on malignant surfaces.

The heterogeneity of surface biomarkers in CRC patients supports a personalized medicine approach to their accurate identification (13). Among identified antigens, Mucin-1 (MUC1) glycoprotein often is overexpressed, relinquishes its apical confinement, and demonstrates abnormal glycosylation patterns (14). Immunohistochemical analysis of MUC1 expression in colon tissue samples from patients showed that abnormal MUC1 expression is present in the earliest stages of colonic tumors and becomes increasingly prominent at later stages of tumor progression (15,16). Our labs previously showed that antibody-functionalized silicon MPs can detect lesions expressing MUC1 in a humanized mouse model (17). Unfortunately, MUC1 is overexpressed on only approximately half of CRC tumors (18,19). Epithelial cell adhesion molecule (EPCAM) is another surface antigen typically expressed at high levels on the basolateral surface of normal epithelial cells (20–22). Overexpression and mislocalization of EPCAM have been linked to cancer progression and unfavorable prognosis in various tumor types (23–29). We developed Antibody-Protein L-MPs (APL-MPs) recognizing MUC1 or EPCAM to test the ability of the multi-antigen approach to achieve higher sensitivity in detecting heterogeneous CRC lesions compared to MPs targeting using either biomarker alone.

## 2. Materials and Methods

### 2.1 Cell culture

The HCT116, SW480, and CACO2 cell lines were purchased from the American Culture Collection (ATCC; Manassas, VA, USA) and maintained in McCoy’s 5A containing 10% (*v/v*) heat-inactivated fetal bovine serum (FBS) (Atlanta Biologicals S11150, Minneapolis, MN, USA), RPMI 1640 (Gibco 22400105, Dublin, Ireland) containing 10% (v/v) FBS, and EMEM (Corning, 10-009-CV, Glendale, AZ, USA) containing 20% (v/v) FBS, respectively. The HT29-MTX-E12 cell line was purchased from Millipore Sigma (St. Louis, MO, USA) and was maintained in DMEM (Corning, 10-013-CV) supplemented with 10% (v/v) FBS and 1% (v/v) non-essential amino acid solution (Gibco, 11140050). The HT29-Luciferase cell line was purchased from Signosis (Santa Clara, CA, USA) and maintained in RPMI 1640 containing 10% (v/v) FBS. All culture media contained 1% (v/v) penicillin/streptomycin (P/S) solution (Gibco 15140122). All cells were passaged at 70-80% confluency with 0.25% (*w/v*) trypsin EDTA (Gibco, 25200072) and incubated at 37°C in a humidified air: CO2 (95:5, v/v) atmosphere.

### 2.2 Immunohistochemical (IHC) staining of human CRC tissues

FFPE human tissue microarrays (TMAs) were acquired from US Biolab (CO2081A, Rockville, MD, USA). Standard IHC protocol was used to stain TMAs for MUC1 and EPCAM antibodies. Briefly, slides were deparaffinized with three 5-minute xylene washes (VWR, 1330-20-7, Missouri City, TX, USA), followed by rehydration through a graded ethanol series(Fisher Scientific, 64-17-5, Waltham, MA, USA): 100% ethanol (2 × 3 min), 95% ethanol (2 × 3 min), 70% ethanol (2 × 3 min), and a final rinse in Milli-Q water (5 min). Heat-induced antigen retrieval with 10 mM sodium citrate (Millipore Sigma, S4641, St Louis, MO, USA) at pH 6 and 95-96°C unmasked the epitopes. The tissue sections were permeabilized with 0.1% (v/v) Triton X-100 in TBST (ThermoFisher, A16046, Waltham, MA, USA) for 10 min at room temperature (RT). Tissue sections were blocked with 10% (v/v) goat serum (Lampire Biological, 7332500, Pipersville, PA, USA) and 1% (w/v) bovine serum albumin (BSA) (Sigma-Aldrich, A9418, St. Louis, MO, USA) in PBS for 60 min at RT and then incubated with primary antibody (1:500) in blocking solution (see above) at 4°C overnight. The next day, sections were treated with 0.3% (v/v) hydrogen peroxide (Millipore Sigma, HX0635, St Louis, MO, USA) diluted in TBS for 10 min at RT, followed by an hour of incubation with sheep anti-mouse HRP-conjugated secondary antibody (Jackson ImmunoResearch, 515-035-062, West Grove, PA, USA) in TBST plus 1% (w/v) BSA for 60 min at RT. After staining with 3,3’-diaminobenzidine (DAB) (Vector Laboratories, SK-4100, Newark, CA, USA) for 10 min at RT, sections were counterstained with Hematoxylin QS for 10 s (Vector, H-3404, Burlingame, CA, USA), dehydrated, and covered with Cytoseal-60 (Epredia, 831016, Kalamazoo, MI, USA) and coverslips. Slides were allowed to dry for at least 1 h before imaging. Slides were imaged using a fluorescence microscope. (Keyence, BZ-X800E; Itasca, IL)

### 2.3 Immunofluorescence (IF) characterization of cell lines in monolayer culture

CRC cell lines were seeded as droplet islands (30,000 cells/droplet) on top of a thin layer of rat tail type I collagen inside a glass-bottom 6-well plate (Cellvis, P06-1.5H-N, Mountain View, CA, USA). Cells were cultured in their corresponding media for 24 h and then fixed with 4% (w/v) paraformaldehyde (PFA). After fixation with 4% (w/v) PFA, cells were stained using a standard IF staining protocol(30). The following primary antibodies and dilutions were used: mouse antibody to MUC1 (Millipore Sigma, 05-652, 1:500), mouse antibody to EPCAM (Santa Cruz Biotechnology, sc-59906, 1:200), and mouse antibody to CEA (Santa Cruz Biotechnology, CI-P-83-1, 1:200). DAPI (ThermoFisher, D3571, Waltham, MA, USA, 1:500) was added to all groups.

### 2.4 Protein L ELISA

Protein L (ThermoFisher, 21189, Waltham, MA, USA) was diluted in carbonate-bicarbonate coating buffer to seven concentrations (0.01, 0.02, 0.05, 0.1, 0.2, 0.5, 1µg/mL). 100 µL of each dilution was added to individual wells of a 96 well plate, the plate was incubated overnight at 4° C, the solutions were aspirated, and the treated wells then were washed three times with a 0.05% (v/v) Tween-20 wash buffer. Non-specific binding sites were blocked with a 1% (w/v) BSA in PBS buffer for 1 h at RT. A non-targeting control (NT-control) mouse IgG antibody (Santa Cruz Biotechnology, sc-2025, Dallas, TX, USA) was added to each well (1:100) and incubated for 1 h at RT. After washing, HRP sheep anti-mouse IgG secondary antibody (Jackson ImmunoResearch, 515-035-062, West Grove, PA, USA) was added (1:5000) and incubated for 1 h at RT. After washing, equal parts of the SuperSignal^™^ ELISA Femto Luminol/Enhancer and SuperSignal ELISA Femto Stable Peroxide Solution (ThermoFisher, 37075, Waltham, MA, USA) were added to each well and mixed for 1 min using a microplate mixer. Relative light units were measured at 425 nm 1-5 mins after adding the substrate using a Cytation 5 Cell Imaging Multi-Mode Reader (BioTek, Winooski, VT, USA).

### 2.5 Primary antibody conjugation and quality control

We used a site-specific antibody labeling kit to label MUC1 antibody with Alexa Fluor^™^ 647 (AF647) and EPCAM antibody with Alexa Fluor^™^ 555 (AF555). Specifically, the SiteClick Antibody Azido Modification Kit (ThermoFisher, S10900, Waltham, MA, USA) was used to modify the carbohydrates in the Fc region of unlabeled IgG antibodies and replace them with an azide-containing sugar. Subsequently, the azide-modified antibody was covalently attached to a Click-iT^™^ sDIBO Alkyne (ThermoFisher, C20028/C20029, Waltham, MA, USA) with fluorescent labels in a copper-free click reaction.

### 2.6 Antibody functionalization of Protein L coated MPs

For *in vitro* studies, MPs were functionalized with primary mouse antibodies and anti-mouse secondary antibodies. For animal studies, MPs were functionalized with fluorescently labeled primary antibodies. Protein L magnetic MPs (ThermoFisher, 88849, Waltham, MA, USA) were washed with 1 mL of 0.05% (v/v) Tween-20 detergent (wash/binding buffer). Primary antibodies were then added to 50 µg of magnetic MPs suspended in 1 mL of wash buffer to achieve a final concentration of 110 µg IgG antibody per mg of MPs. The MPs were then incubated at RT for 1 h. After the incubation period, the MPs were washed twice with wash buffer, separating the MPs from the buffer using a magnet following each wash step. For *in vitro* studies, goat-anti-mouse IgG-Alexa Fluor^™^ 488 (AF488) secondary antibody (ThermoFisher, A11001, Waltham, MA, USA) was added (1:1000) to the Protein L-MPs in 1 mL of wash buffer and incubated at RT on a rotary mixer for 1 h. After incubation, MPs were washed three times with wash buffer for 5 min at RT. After the 3rd wash, MPs were suspended in culture media and incubated with cell tumoroids.

### 2.7 Tumoroid formation assay

For IF staining and *in vitro* binding experiments, HT29-MTX, CACO2, or HT29-Luciferase cell lines were seeded at a density of 100,000 cells per well into 24-well plates containing 12 wells with 750 round-bottomed microwells (5D spherical plate, Kugelmeires, Zurich, Switzerland). Microwell plates were placed on a rotary shaker at 60 rpm for 48 h. Tumoroids were harvested using a wide bore 1000 µL pipette tip and transferred to a 1.5 µL tube for live binding assay or fixed with 4% (w/v) PFA for IF staining.

### 2.8 Binding of antibody functionalized MPs to cell tumoroids

Protein L-MPs functionalized with mouse IgG antibodies targeting either MUC1 or EPCAM, or with NT-control mouse IgG antibody, were suspended in 500 µL of culture media prior to the binding assay. HT29-MTX or HT29-Luciferase tumoroids suspended in 1 mL of culture medium were added to each group of APL-MPs and incubated at 37°C on a rotary shaker at 60 rpm for 1 h. After the incubation period, the tumoroid/MP suspension was transferred to a 100 µm cell strainer and washed thrice with 50 µL of PBS. Live tumoroids then were transferred to a glass bottom six-well plate containing 1 mL of culture media and imaged with a Nikon A1R-MP confocal microscope (Nikon Corporation, Japan).

### 2.9 Co-culture of HT29-MTX and HT29-Luciferase cells in mixed monolayers and as HET-tumoroids

HT29-MTX and HT29-Luciferase cells were co-cultured as monolayers and as tumoroids, at different mixing ratios. For 2D co-cultures, HT29-MTX/HT29-Luciferase cells were seeded in a T-75 culture flask (Corning, 07-202-00) at the following ratios: 50/50, 75/25, 25/75, 100/0, and 0/100. The cells were cultured for 3 and 10 days as two separate groups. The cells in culture underwent routine change of media containing RPMI 1640 and DMEM plus 10% (v/v) FBS at a 50/50 ratio and were passaged at 80% confluency. After 3 and 10 days of co-culture, the cells were seeded in a glass-bottom 6 well pate for 48 h before fixation with 4% (w/v) PFA and staining with rabbit anti-luciferase (1:500, Millipore Sigma, L0159, St. Louis, MO, USA), MUC1 (1:500) primary antibodies, and DAPI (1:500), goat anti-rabbit Alexa Fluor^™^ 568, and mouse anti-human AF488 secondary antibodies (ThermoFisher, A11036, A11001, Waltham, MA, USA). The plates then were imaged with a Nikon A1R-MP microscope.

To create HET-tumoroids, HT29-MTX and HT29-Luciferase cells were first tagged with fluorescent MitoTracker Deep Red MF and MitoTracker Green MF (ThermoFisher, M22426, M7514, Waltham, MA, USA), respectively. Briefly, lyophilized MitoTracker was dissolved in anhydrous dimethyl sulfoxide (DMSO) to prepare a stock solution with a final concentration of 1 mM. The optimal concentrations for staining live cells were determined to be 120 nM and 300 nM for MitoTracker Green MF and MitoTracker Deep Red MF, respectively. HT29-MTX and HT29-Luciferase cells were suspended in 100 µL of pre-warmed (37°C) MitoTracker staining solution and growth media for 45 min. After staining, the cells were washed with growth medium, centrifuged, resuspended in growth medium, and co-cultured in a microwell plate at the following mixing ratios (HT29-MTX/HT29-Luciferase): 90/10, 80/20, 70/30, 60/40, 50/50, 40/60, 30/70, 20/80, 10/90, 100/0, and 0/100. The co-culture was maintained in a 1:1 (v/v) mixture of RPMI 1640 and DMEM for 72 h before live imaging inside a glass-bottom six-well plate.

### 2.10 Mouse flank tumors to test heterogeneous tumor induction and detection by APL-MPs

All animal experiments were conducted in accordance with international standards. All animal experimental procedures were approved by the University of Texas MD Anderson Institutional Care and Use Committee (Protocol 1801, Approved 7/27/2021). In two initial pilot experiments, we injected 100,000 HT29-MTX and HT29-Luciferase single cells suspended in 100 µL PBS at three different mixing ratios (HT29-MTX/HT29-Luciferase): 75/25, 50/50, and 25/50 into the flanks of 9-week-old female nude mice (Envigo/Harlan Labs, Indianapolis, IN, USA, Athymic nude (nu/nu), Stock number 069). Tumor size and bioluminescence signal were monitored using calipers and an IVIS Spectrum CT small animal imager (PerkinElmer, Waltham, MA, USA) to determine the optimal ratio of CRC cells and the preferred mode of injection (single cell vs. tumoroids).

Based on these results, we next injected 1 million HT29-MTX/HT29-Luciferase cells, suspended in 200 µL of either PBS or a 50/50 (v/v) solution of PBS + Matrigel^™^, into the left and right flanks of four 9-week-old female nude mice. Injections on the left side contained Matrigel™, whereas injections on the right side did not. Tumor size and bioluminescence signal were monitored using calipers and an IVIS Spectrum CT small animal imager, as mentioned previously, to evaluate the impact of Matrigel^™^ on tumor size and bioluminescence signal. The flank tumors were removed approximately 4 weeks after injection after they reached 10 mm in diameter. Tumors then were sliced with a #10 sterile surgical blade (Fisher Scientific, 22-079-683, Waltham, MA, USA) and were incubated with MUC1-, EPCAM-, and a 50/50 (w/w) combination of the two singly-labeled species (“MUC1 + EPCAM”)- Protein L functionalized MPs for 1 hr at 37°C. The tumor sections were then fixed with 4% (w/v) PFA for 12 h, stained with 5 µg/mL of DAPI in PBS, and imaged with a Nikon A1R-MP confocal microscope using a 20X objective lens.

### 2.11 Colonic submucosal injection of CRC cell lines in mice to create HET-tumors

HT29-MTX and HT29-Luciferase cell lines were prepared for orthotopic injection into the colon of 5-to 9-week-old female nude mice (Envigo/Harlan Labs, Indianapolis, IN, USA, Athymic Nude (nu/nu), Stock number 069) as single cells and tumoroids in two separate groups. The first group of mice was injected with 500,000 HT29-MTX/HT29-Luciferase single cells at 30/70 ratio (30% HT29-Luciferase and 70% HT29-MTX) to create HET-tumors expressing MUC1 and EPCAM tumors. The second group of mice was injected solely with 500,000 HT29-Luciferase single cells (MUC1^−^/EPCAM^+^). The final injection volume was 50 µL in serum-free RPMI 1640 media, including 65% cells (32.5 µL), 25% (12.5 µL) Matrigel^™^, and 10% (5 µL) methylene blue (Neopharma, Aschau, Germany). For tumoroid injections, HT29-MTX and HT29-Luciferase tumoroids were cultured separately in microwell plates as described above, but with 25,000 cells per well instead of 100,000 cells. The lower seeding density resulted in the formation of smaller tumoroids (approximately 50-60 µm in diameter) that would travel undisturbed through a 33 G needle (inner diameter 108 µm Hamilton, 7803-05). Tumoroids with diameters greater than 70 µm were removed using a 70 µm cell strainer. The filtered tumoroids were centrifuged at 700 RPM for 1 min, and the pellets were resuspended in 500 mL serum-free RPMI 1640 medium. HT29-MTX and HT29-Luciferase tumoroids were suspended in serum-free media and then mixed at 30/70 ratio (30% HT29-Luciferase and 70% HT29-MTX). The resulting mixture was centrifuged again at 700 RPM for 1 min. The pellet was resuspended in 45 µL of serum-free RPMI 1640 medium plus 5 µL of ink and injected orthotopically into the colon of nude mice. Matrigel^™^ was not added to the tumoroid injections.

Colonic submucosal injection was performed using a Coloview mini endoscope system (Vetcom; Karl Storz, Tuttlingen, Germany) with Endovision Tricam (Karl Storz), Hamilton 50 µL Microliter Syringe (Hamilton, 80530, Reno, NV, USA), and a 10-inch, 30-degree, 33-gauge, beveled Hamilton Small Hub removable needle (Hamilton, 7803-05, Reno, NV, USA), according to a previously described method (31). Prior to colonic submucosal injection, the microliter syringe was filled with 50 µL of cells, Matrigel^™^ and ink mixture and kept on ice to avoid gelation at RT. Submucosal injection was performed through the working channel of the endoscope. The flexible 33-guage needle was inserted through the working channel of the endoscope. With the bevel facing up, the needle was gently introduced into the mucosa > 1 cm proximal to the rectal opening. Cancer cells mixed with Matrigel^™^ and ink then were injected by a second investigator. Mucosal lifting confirmed a successful injection (Video SI.1).

### 2.12 Bioluminescence imaging of mice after submucosal injection

After submucosal injections, bioluminescence (BLI) imaging was performed weekly using an IVIS Spectrum CT small animal imager (PerkinElmer, Waltham, MA, USA). Mice were anesthetized with isoflurane and received 200 µL (15 mg/mL in DPBS) of D-luciferin and potassium salt (GoldBio, LUCK-1G, St. Louis, MO, USA) via intraperitoneal (i.p.) injection. BLI images were captured 10 min after injection with mice in the supine position. Living Image Software was used in auto-exposure mode with medium binning, F/Stop of 1, and Field of View set to D.

### 2.13 In vivo MP binding experiments

To investigate the specific binding of APL-MPs to HET-tumoroids *in vivo*, a two-arm animal study involving 54 mice was conducted. Each group was divided into subgroups that received MUC1, EPCAM, MUC1 + EPCAM, or NT-control mouse IgG APL-MPs. Prior to the *in vivo* MP binding studies, the animals were anesthetized with isoflurane and kept under anesthesia for the duration of the experiment. The mouse colon first was flushed with 1 mL of PBS three times using a gavage needle (VWR, 89233-486, Missouri City, TX, USA). Subsequently, 50 µg of APL-MPs suspended in 500 µL of Hanks’ balanced salt solution (HBSS) (ThermoFisher, 14025092, Waltham, MA, USA) was administered through the rectum. After a 30 min incubation period, each animal was sacrificed and the colon was removed. The colon was then further flushed with HBSS, opened along its length, and mounted on 1/4 inch poly(dimethylsiloxane) (PDMS) (Dow Corning, Sylgard 184, Midland, MI, USA) inside a 6-well plate. The colon then was fixed with 4% (v/v) PFA for 12 h at 4°C. After fixation and removal of PFA, the colon sections were washed with PBS and permeabilized with 0.5% (v/v) Triton-X100 for 2 h. Following permeabilization, the colon sections were incubated with a rabbit-anti-human nuclear antigen (HNA) antibody (NeoBiotechnologies, RBM5-346-P1, Union City, CA, USA) at a final concentration of 1 µg/mL at RT for 1 h. The colon then was washed and incubated with 5 µg/mL DAPI and goat-anti-rabbit IgG-AF488 secondary antibody (ThermoFisher, A-11034, Waltham, MA, USA) at a 1:1000 dilution ratio for 1 h at RT. The colon sections were imaged inside a glass-bottom six-well plate. To stabilize the colon during confocal imaging, a PDMS mold was created from the wells of a 6-well plate. The mold then was positioned within the well containing the colon tissue sample. The mold was secured tightly in the well by enhancing the friction on its sides using laboratory tape.

### 2.14 Confocal imaging and imaging of the mouse colon

IF imaging of the mouse colon was conducted in an automated fashion using a Nikon A1R-MP confocal microscope. The acquisition process was streamlined with the aid of the NIS-Elements JOBS module, capturing three sets of nine (3 × 3 grid) images at random regions of the colon using a 20X objective, with approximately 40 µm Z-stacks at 0.89 µm steps.

### 2.15 Image processing and quantification of particle binding

Quantification of MP binding was performed using IMARIS image analysis software (Bitplane, Zurich, Switzerland) and custom code written in MATLAB (MathWorks, Natick, MA, USA). Stitched z-stack images with constant length along the x- and y-axes and variable depth along the z-axis were imported into the IMARIS software. The image dimensions spanned 2868 pixels in both the x- and y-axes, and less than 100 pixels in the Z-axis. The respective voxel sizes were 0.621, 0.621, and 0.875 µm in the x, y, and z directions. The Spots function was used to identify the mouse colon epithelial cell nuclei (DAPI), nuclei of human cancer cells (HNA/DAPI), and MPs based on the fluorescent signals in each channel throughout the entire z-stack. Expected spot diameters of 5 µm, 7 µm, and 10 µm were defined for DAPI, HNA, and MPs, respectively. The spot size was estimated from the image and kept the same for all experiments. It is worth noting that the estimated diameter did not impact the quantification process, as the spot center-to-center distance was used in the analysis rather than their boundaries. To eliminate background noise, a Gaussian filter was applied, effectively smoothing objects located within a distance less than 8/9 of the anticipated spot radius. Spot quality thresholds were determined manually and separately for each channel to account for variations in fluorophore signal intensity in each experiment. These thresholds were set just below the point at which a distinct increase in background false positives was observed. The resulting x-, y-, and z-position coordinates of all spots were imported into MATLAB for further analysis.

To determine the sensitivity and specificity of tumor detection, we partitioned the 3D image into a grid of equidistant boxes along the x and y dimensions, with only one box along the z dimension. In one such instance, the grid consisted of 24 boxes in the x-direction and 24 in the y-direction, thereby creating 576 unique boxes. This grid-based analysis allowed us to classify spatially the presence of MP objects in the same grid box as the HNA^+^ objects, with these identifiers: true positive (TP), false positive (FP), true negative (TN), and false negative (FN). The classification criteria were as follows:

- **TP**: The grid box contained at least one object from the HNA channel and one object from the MP(s) channel.
- **FP**: The grid box contained at least one object from the MP(s) channel but none from the HNA channel.
- **FN**: The grid box contained at least one object from the HNA channel but none from the MP(s) channel.
- **TN**: The grid box is devoid of objects from both the HNA and MP(s) channels.

The sensitivity and specificity were calculated using the following formulas:

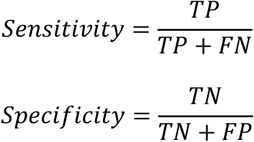

To ensure the accuracy of the specificity calculations, we required the presence of at least one DAPI object within a grid box to consider it for specificity calculations. This approach helped eliminate potential false positives and provided a more reliable estimation of test specificity.

### 2.15 Statistical Analysis

One-way analysis of variance (ANOVA) was conducted using GraphPad Prism (GraphPad Software version 7.03, San Diego, CA, USA) to compare the detection sensitivity and specificity among four types of APL-MPs: MUC1, EPCAM, MUC1 + EPCAM, and NT-control mouse IgG1. The ANOVA included sensitivity and specificity values for each group, with sample sizes ranging from n = 3 to n = 4. Tukey’s secondary multiple comparisons test was used to evaluate significant differences between group means.

### 2.16 Data availability

Raw data are available from the corresponding author upon request for research or academic purposes.

## 3. Results

### 3.1. Expression of MUC1 and EPCAM in CRC cell lines reflect cancer heterogeneity

We first investigated the expression of cell-surface antigens EPCAM and MUC1 in four CRC cell lines (CACO2, HT29-MTX, HCT116, and SW480). Antigens were selected based on their heterogeneous expression levels in human CRC tissue sections (representative sections are shown in **Fig. 1A**). IF staining of CRC cell lines in 2D monolayers revealed different patterns of antigen expression, consistent with their heterogeneous tumor origins. As shown in **Fig. 1B**, MUC1 was highly expressed in HT29-MTX and SW480 cells (MUC1^hi^), but was weakly expressed in CACO2 and HCT116 cells (MUC1^low^). EPCAM was expressed strongly in all cells. Given the higher expression of MUC1 and EPCAM in HT29-MTX cells and weak MUC1 expression in CACO2 cell lines, HT29-MTX and CACO2 cells and MUC1 and EPCAM targets were selected for subsequent studies.

**Figure 1.**
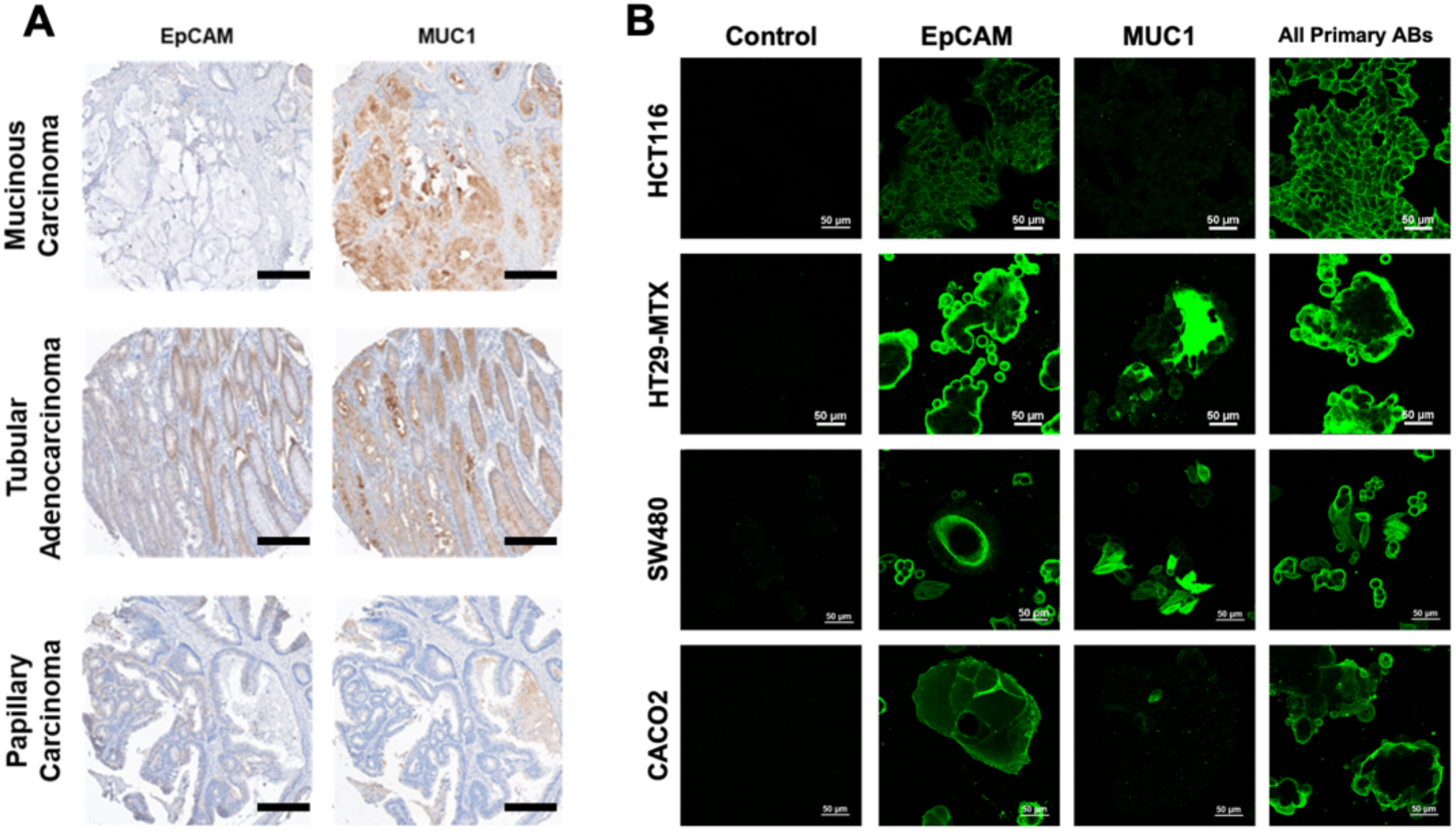
IHC and IF staining show that surface markers MUC1, EPCAM are heterogeneously expressed. (**A**) IHC staining (brown = positive) for EPCAM and MUC1 in mucinous adenocarcinoma, tubular adenocarcinoma, and papillary carcinoma human tissue samples. Scale bar = 0.2 mm. (**B**) IF staining (green = positive) for EPCAM and MUC1 in CRC cell lines, revealed ubiquitous EPCAM expression across all cell lines and high MUC1 expression only in SW480 and HT29-MTX cells.

### 3.2 Tumoroids preserve antigen expression from 2D to 3D and mixed tumoroid models recreate heterogeneous antigen expression in defined ratios

We investigated the expression patterns of surface antigens MUC1 and EPCAM in 2D and 3D organoids. The HT29 variants HT29-MTX and HT29-Luciferase showed the most complementary expression of these markers in 2D cultures. To investigate marker expression in 3D, we formed tumoroids of each, using low-adherence culture plates (**Fig. 2A**), and then immunostained each (**Fig. 2B**). We observed high MUC1 expression on HT29-MTX tumoroids and weak sporadic staining (MUC1^low^) on HT29-Luciferase tumoroids. Conversely, both HT29-MTX and HT29-Luciferase tumoroids expressed EPCAM (EPCAM^hi^); however, the expression pattern of EPCAM on HT29-MTX tumoroids was notably irregular and less organized than that on HT29-Luciferase tumoroids.

**Figure 2.**
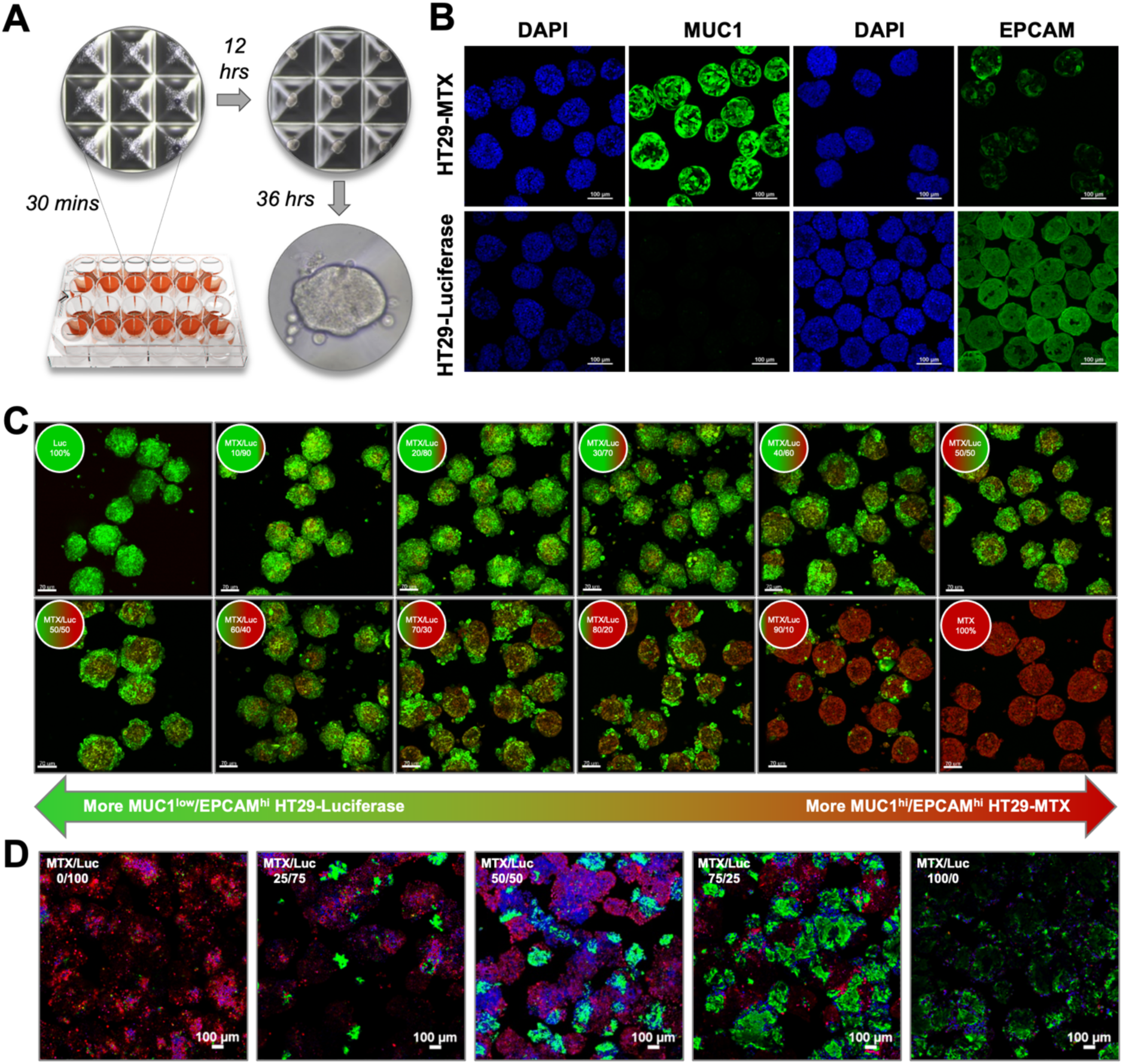
Signature MUC1 and EPCAM expression patterns in HT29-MTX and HT29-Luciferase tumoroids enables culture as mixed HET-tumoroids. (**A**) Schematic of clustering plate with brightfield images of HT29-MTX cells 30 min, 12 hrs, and 36 hrs after seeding cells. **(B)** IF staining of HT29-MTX and HT29-Luciferase tumoroids for MUC1 and EPCAM. (**C**) **A**) *In vitro* co-culture of labeled HT29-Luciferase (Luc, green) and HT29-MTX (MTX, red) cell lines as HET-tumoroids at various mixing ratios. Cell line on the exterior surface of tumoroids was evaluated by live imaging after 3 days of *in vitro* culture. Scale bar = 70 µm. (**D**) Expression of MUC1 and EPCAM was assessed in 2D co-culture mixtures of HT29-Luciferase and HT29-MTX cells at different mixing ratios through IF staining, revealing distinct patterns of EPCAM (red) and MUC1 (green) expression. All confocal microscopy images in (**B-D**) are maximum intensity projections from a multilayer z-stack.

We next sought to create representative heterogeneous tumoroids (HET-tumoroids) built from mixtures of HT29-MTX and HT29-Luciferase cells (**Fig. 2C**). Labeled HT29-MTX and HT29-Luciferase cells were co-clustered as HET-tumoroids at different mixing ratios, aiming to identify an optimal ratio that enabled equal surface accessibility for each cell type and its signature antigens. As shown in **Fig. 2C**, live imaging 72 h after co-culture revealed a consistent presence of both cell lines on the exterior of HET-tumoroids in the 70/30 ratio group (HT29-MTX/HT29-Luciferase). IF staining of a subset of these HET mixtures (**Fig. 2D**) demonstrated that both MUC1 and EPCAM antigens were accessible by antibodies in combination co-cultures, particularly at a 75/25 ratio, similar to the data in **Fig. 2C**. EPCAM expression in HET-tumoroids with ≥ 70 % HT29-MTX had high, but non-uniform, staining similar to monocultures in Fig. 2B.

### 3.3. IgG Antibodies bind to Protein L and APL-MPs can detect CRC surface antigens in tumoroids and in tissue

We employed IgG-binding Protein L as an intermediate adapter to functionalize micron-sized iron oxide magnetic particles (1 µm nominal diameter) with IgG1 antibodies for the detection of CRC surface antigens (**Fig. 3A**). As a first step, we evaluated the binding capacity of surface-bound Protein L to a non-targeting control (NT-control) mouse IgG1 antibody using an enzyme-linked immunosorbent assay (ELISA) (**Fig. SI.2**). As expected, Protein L exhibited strong binding capacity to the NT-control mouse IgG1 antibody, with negligible background binding of fluorophore-labeled goat IgG1 secondary antibody (32). We exposed MPs, pre-coated with Protein L, to labeled IgG1 control and confirmed that binding saturation was reached at 110 µg/mg of ms IgG1 for MPs of this size, specific surface area, and surface coverage with Protein L (**Fig. SI.3)**.

**Figure 3.**
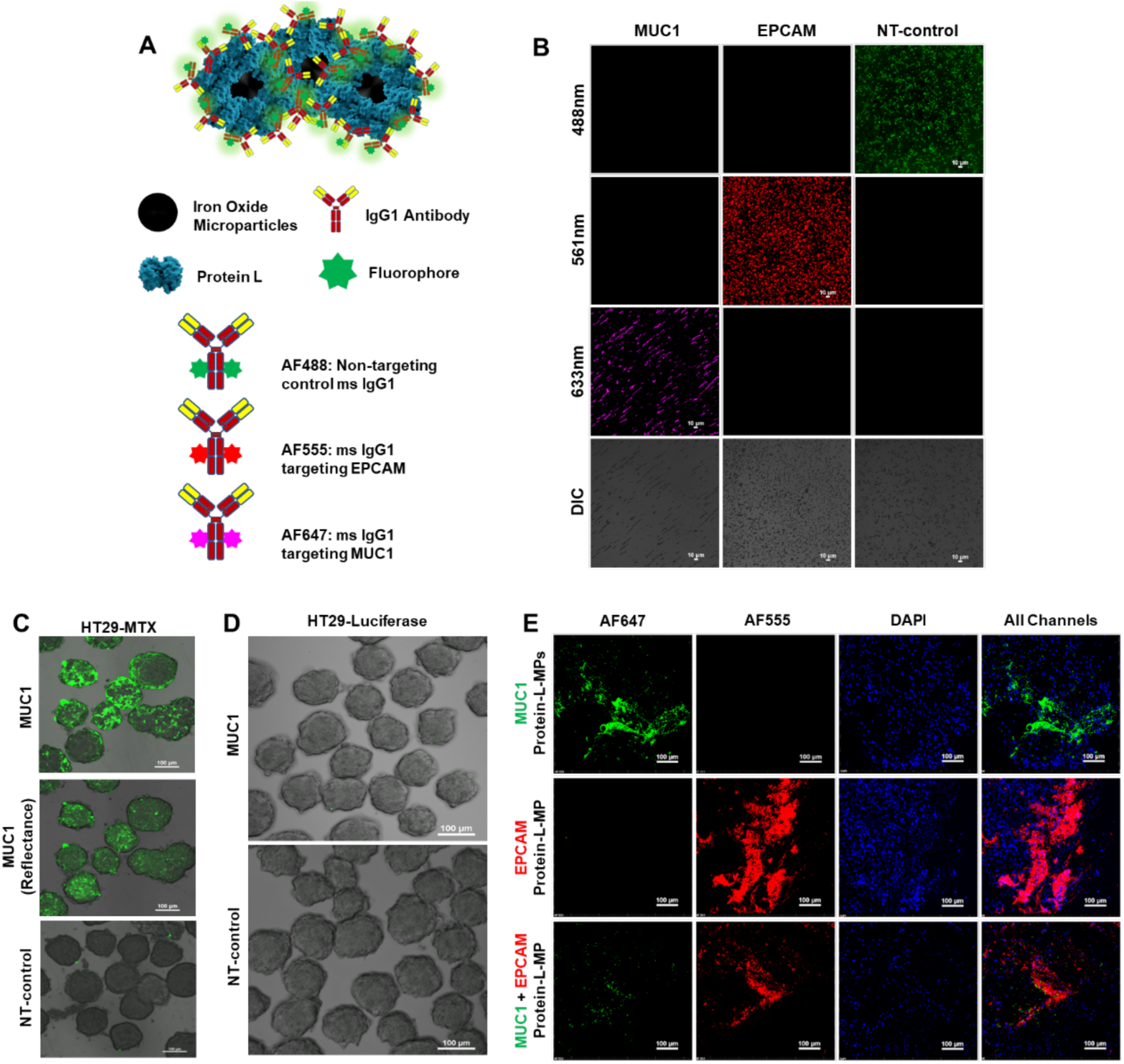
APL-MPs recognize human antigens on CRC cells, tumoroids, and tissues. (**A**) Schematic of Protein L-MPs functionalized with Fc-labeled ms IgG1 antibodies (**B**) Fluorescent imaging of APL-MPs confirming antibody presence on each MP. (**C**) Specific binding of MUC1 APL-MPs to HT29-MTX tumoroids imaged in fluorescent mode and confirmed with reflectance imaging. NT-control ms IgG1-Protein L-MPs did not bind to HT29-MTX tumoroids. (**D**) MUC1 APL-MPs and control IgG1-protein-MPs did not bind to MUC1^low^ HT29-Luciferase tumoroids. (**E**) Tissue sections from flank HET-tumors were incubated with MPs targeting either MUC1, EPCAM, or a combination of both MPs. Specific binding of the MPs to the HET-tumoroids was observed, with non-overlapping signals in the overlaid image.

For antigen binding experiments, we used the same Protein L-coated MPs, functionalized with either mouse IgG1 antibodies against MUC1 or EPCAM, or the NT-control (**Fig. 3A**). Covalent labeling of these antibodies with Alexa Fluor™ fluorophores via click chemistry attachment to the Fc region enabled visualization of the APL-MP conjugates (**Fig. 3B**), with fluorescence specific to each biomarker. Retention of labeled antibodies was confirmed by increased UV absorbance at 280 nm, reflecting the bound protein content (**Fig. SI.4**).

To investigate the specific binding of APL-MPs to MUC1 and EPCAM antigens on CRC cells, we conducted binding experiments using homogeneous MUC1^hi^/EPCAM^hi^ HT29-MTX and MUC1^low^/EPCAM^hi^ HT29-Luciferase tumoroids. Specific binding of MUC1 APL-MPs was observed exclusively on HT29-MTX tumoroids, while no binding was observed to HT29-Luciferase tumors (**Fig. 3 C-D**). We confirmed specificity using confocal microscopy to visualize the fluorescent signal from surface-tethered antibodies along with a reflectance signal from the MPs. NT-control mouse IgG1 APL-MPs did not bind to either of the homogeneous tumoroids, further confirming the specificity of MUC1 APL-MPs. EPCAM APL-MPs bound to both homogeneous (HT29-MTX and HT29-Luciferase) tumoroids, indicating successful recognition of EPCAM antigens.

A series of experiments were conducted to establish HET-tumoroids *in vivo* from mixtures of HT29-MTX and HT29-Luciferase cells and to confirm their display of accessible surface markers. HT29-MTX and HT29-Luciferase cells were injected subcutaneously as single cell mixtures with or without Matrigel^™^ into the flanks of 9-week-old female nude mice to assess optimal growth at three cell ratios of HT29-MTX/HT29-Luciferase: 75/25, 50/50, or 25/75. Larger tumors were identified by bioluminescence and caliper measurements when single-cell suspensions in PBS and Matrigel^™^ were injected (**Fig. SI.5**). Confocal imaging of sections of flank tumor, after incubation with either MUC1 APL-MPs, EPCAM APL-MPs, or 50:50 mixtures of the two, revealed specific binding to HET-tumors (**Fig. 3E**). In addition, we observed negligible background fluorescence from NT-control IgG1 APL-MPs in the flank tumors (**Fig. SI.6**)

Based on *in vitro* HET-tumoroid studies and *in vivo* flank experiments, a 70/30 ratio of cells (HT29-MTX/HT29-Luciferase) was chosen as optimal for colonic submucosal injection to form orthotopic HET-tumors.

### 3.5. Endoscopy-guided orthotopic implantation of CRC cells enables colonic HET-tumors in mice

To optimize CRC HET-tumor formation, we compared submucosal implantation of 500,000 single cells mixed with Matrigel^™^ and 500,000 cells as tumoroids without Matrigel^™^ into the colons of mice under endoscopy guidance (**Fig. 4A**). Both groups received CRC cells at a 70/30 ratio (HT29-MTX/HT29-Luciferase). Four weeks post-injection, three of four mice that received single-cell mixtures with Matrigel^™^ developed tumors, while only one of three mice with HET-tumoroid injections exhibited tumor formation. Through bioluminescence imaging via D-luciferin injection and endoscopic assessment of the mouse colon, we found that the optimal time to conduct *in vivo* binding experiments was ∼35 days post-implantation of CRC cell mixtures (**Fig. 4B**). Visual examination of colons injected with CRC cell mixtures revealed wall thickening compared to normal mouse colon (**Fig. 4C**). Confocal microscopic imaging of the luminal surface of normal colon tissue and tumor-adjacent normal tissue identified the typical crypt structure observed in healthy colonic tissue, as evidenced by DAPI staining and negative staining for human nuclear antigen (HNA). In contrast, analysis of CRC-injected colons revealed the presence of CRC cells, characterized by disorganized nuclei and abundant HNA signal, confirming the retention and expansion of human cancer cells.

**Figure 4:**
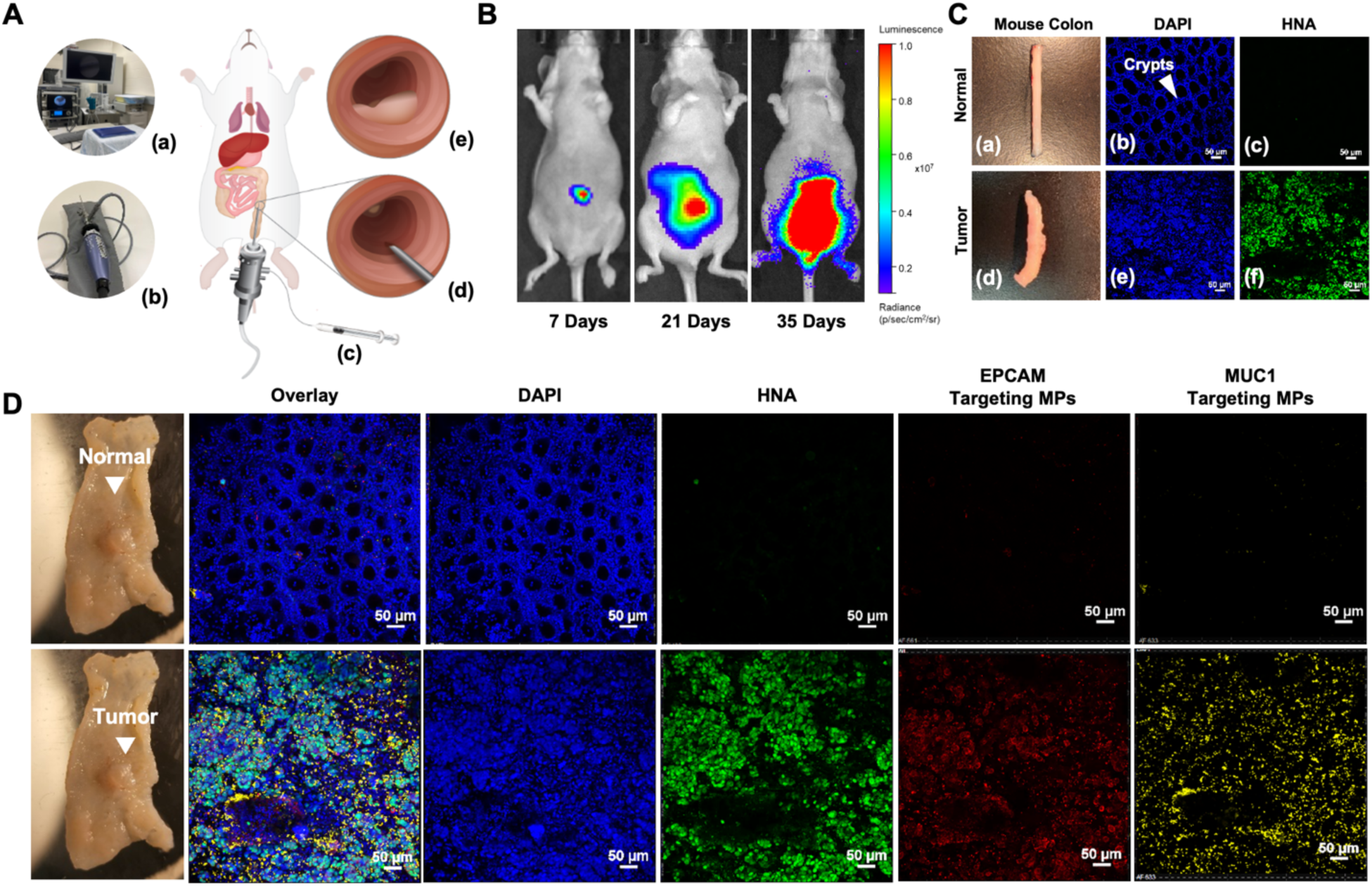
Minimally invasive submucosal implantation of human CRC cell mixtures to generate HET-tumors in an orthotopic mouse model. (**A**) Overview of the procedure with subparts showing photographs of (a) the instrument suite and (b) the mini-endoscopic system; schematics of (c) a microliter syringe and a flexible injection catheter, (d) zoomed view of the injection needle entering the submucosal tissue, and (e) successful implantation indicated by mucosal lifting. **(B)** Bioluminescence imaging of mice at 7 , 21, and 35 days post-implantation demonstrates tumor progression in the colon. (**C**) Visual comparison between (a) normal colon tissue and (d) colon tissue injected with CRC cells. IF imaging of the interior, luminal surface of normal mouse colon tissue with (b) DAPI and (c) HNA demonstrated normal colonic crypt structures and the absence of CRC cells. In contrast, the colon tissue with HET-tumor displayed (d) thickened walls, (e) unorganized nuclear staining, and (f) positive HNA signal confirming the presence of CRC cells. (**D**) Representative images of areas of normal mouse colon tissue, and areas of human CRC tumors. After *in vivo* incubation with targeting MPs, colons were excised, rinsed, fixed, and IF-stained for HNA. Confocal imaging of the luminal side showed normal tissue organization into crypts, with minimal signal in other detection channels. Tumor-bearing regions displayed aberrant organization of cell nuclei (DAPI channel), corresponding to visible overgrowth of human CRC cells (HNA channel). Both EPCAM-targeting and MUC1-targeting MPs identified HNA^+^ regions.

### 3.6. APL-MPs detect CRC surface antigens on the luminal colon surface in vivo

To investigate specific binding of MUC1 APL-MPs and/or EPCAM APL-MPs to HET tumors *in vivo*, we designed a two-arm animal study of 54 mice. In the first group, 27 mice received orthotopic implantation of 70%/30% MUC1^hi^/EPCAM^hi^ HT29-MTX and MUC1^low^/EPCAM^hi^ HT29-Luciferase cell mixtures submucosally to create HET-tumors. The second group of 27 mice received orthotopic implantation of MUC1^low^/EPCAM^hi^ HT29-Luciferase cells to form homogeneous EPCAM^hi^ tumors. This group served as a control to investigate the effects of multi-antigen vs. single-antigen targeting. Of the total of 54 mice injected, 31 mice (56%) developed tumors, and 26 of these mice were reserved for binding experiments, confocal imaging, and subsequent quantification of particle binding. Upon tumor maturation, anesthetized mice in each group received an intracolonic suspension of APL-MPs in HBSS, functionalized with either MUC1, EPCAM, or control IgG1, or a 1:1 combination of singly functionalized MUC1 APL-MPs and EPCAM APL-MPs. Following a 30 min incubation with APL-MPs *in vivo,* colons were removed, rinsed with HBSS, formaldehyde-fixed, bisected longitudinally, stained with DAPI and HNA as before, and imaged on the luminal face by confocal microscopy (**Fig. 4D**). Imaging confirmed the expected normal tissue morphology in non-injected regions: colonic crypts with no detectable HNA signal and no detectable retention of MUC1-targeting or EPCAM-targeting APL-MPs. Regions with macroscopic tumors contained tissue-level changes: crypt structures were lost, overrun by human CRC cells, and fluorescent MUC1-targeting and EPCAM-targeting APL-MPs identified MUC1^hi^ and EPCAM^hi^ HET tumors, respectively. Negligible binding of control IgG1 APL-MPs to tumors was observed.

### 3.7. Targeting multiple surface antigens enhances heterogenous HET-tumor detection sensitivity

To quantify the sensitivity and specificity of detection of individual and combined APL-MPs, a grid-based analysis approach was used to analyze tissue images (**Fig. 5A**). This approach involved segmenting signals for each individual nucleus, each HNA^+^ nucleus, and each labeled MP. The positional relationships between all segmented objects, both in relation to one another and with respect to the grid box, were examined. The MP-based identification of CRC, as validated by an adjacent HNA signal, and appropriate absence of MPs from non-tumoroid regions were identified as true positive (TP) and true negative (TN), respectively. Misidentification of tumor and normal tissues were identified as false positive (FP) and false negative (FN) (**Fig. 5A (m))**. By systematically varying the grid size (**Fig. 5A (j-l))**, we captured the sensitivity and specificity of APL-MPs at various detection resolutions. **Fig. 5B** shows example tissues with APL-MP detection for each biomarker, overlaid with nuclei and HNA signals, and the resulting graphical analysis on a 24x24 grid. Additional qualitative examples of these grid analyses are presented in **Fig. SI.7**. Quantified analysis of MP sensitivity to tumor detection (defined in Methods) showed comparable results for MUC1 and EPCAM detection of HET-tumors, but significantly higher sensitivity when both MUC1-targeting and EPCAM-targeting APL-MPs were used concurrently (**Fig. 5C ).** All were higher than the baseline NT-control IgG. For tumors comprising only HT29-Luciferase cells (MUC1^low^/EPCAM^hi^), MUC1 APL-MPs did not improve sensitivity above the IgG control baseline, nor did they improve detection above EPCAM APL-MPs. These data showed that the CRC biomarkers were as expected, accessible to the APL-MPs, and identified by the APL-MPs. They also supported our hypothesis that the combination of APL-MP types would improve tumor identification.

**Figure 5:**
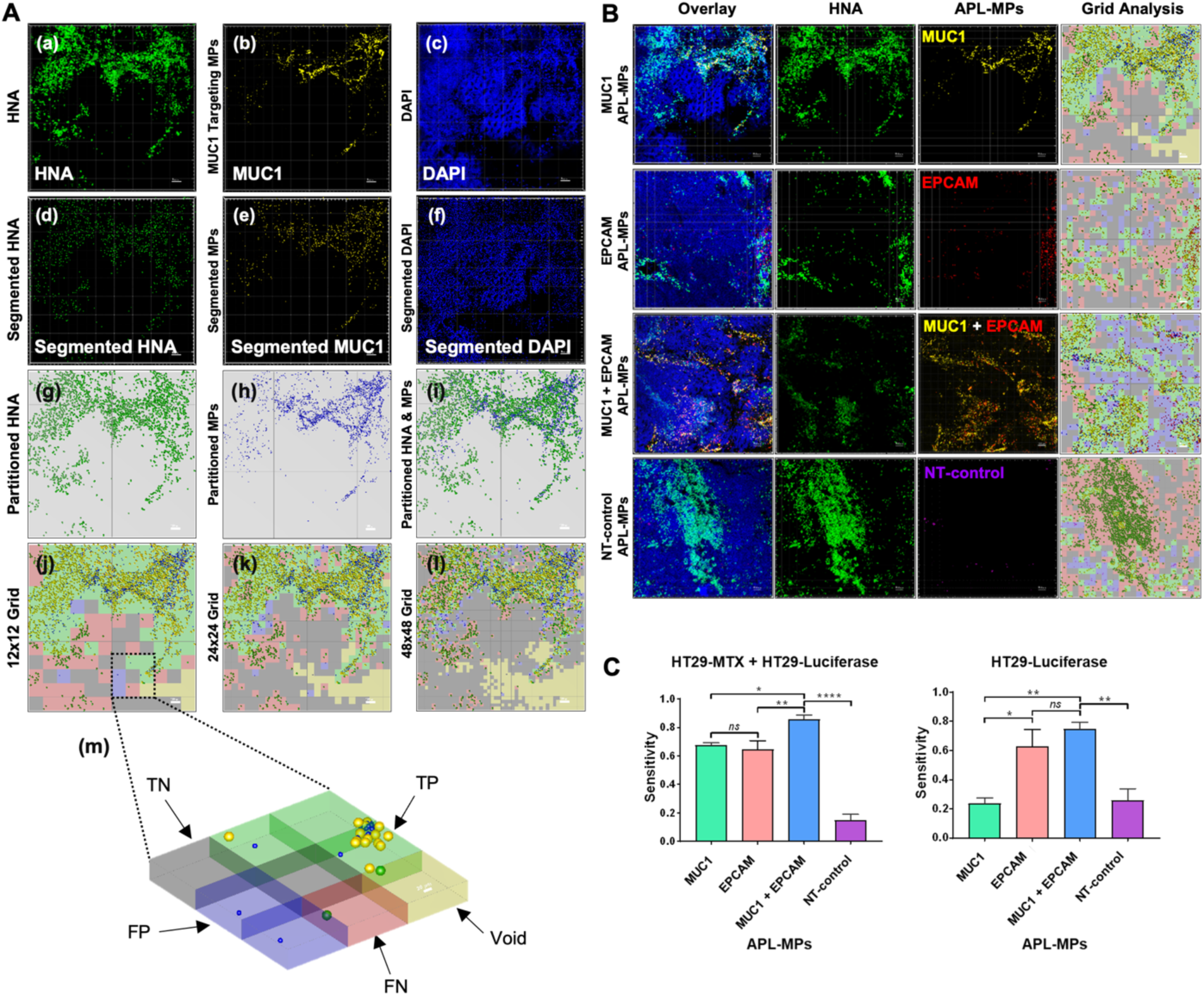
Colonic tissues incubated with APL-MPs *in vivo* demonstrate specific antigen targeting. (**A**) Development of a grid-based approach to evaluate sensitivity and specificity of individual and combined APL-MPs. (a-f) Confocal imaging and segmentation: (a) HNA channel (green), (b) MUC1 APL-MPs (yellow), (c) DAPI channel (blue), (d) HNA segmentation, (e) MUC1 APL-MPs segmentation, and (f) DAPI segmentation. (g-l) Grid-based analysis: (g) Spheres representing HNA on a 12x12 grid, (h) Spheres representing MUC1 APL-MPs on a 12x12 grid, (i) Overlay of HNA and MUC1 spheres on a 12x12 grid, (j) Analyzed 12x12 grid highlighting TP, TN, FN, FP, and void regions, (k) Regional analysis using a 24x24 grid, (l) Regional analysis using a 48x48 grid, and (m) Zoomed-out view of 6 grid boxes emphasizing TP, TN, FN, FP, and voids. (**B**) Confocal images of HT29-MTX/HT29-Luciferase HET-tumors being detected with MUC1, EPCAM, and MUC1+EPCAM APL-MPs. Control NT-control IgG1 APL-MPs exhibited minimal to no binding to CRC cells, indicating their lack of specific interaction. Cancer cell nuclei (HNA) shown in green, MUC1 APL-MPs shown in yellow, EPCAM APL-MPs shown in red, control IgG1 APL-MPs shown in purple. Analyzed 24x24 grids highlighting areas of TP, TN, FP, FN is shown on the right. (**C**) Sensitivity analysis: Comparison of individual and combined APL-MPs targeting MUC1^hi^/EPCAM^hi^ HT29-MTX/HT29-Luciferase HET-tumors (left) and MUC1^low^/EPCAM^hi^ HT29-Luciferase homogeneous tumors (right), using a 24x24 grid (75 µm detection resolution). * p < 0.05, ** p < 0.01, *** p < 0.001, *ns* (not significant). One-way ANOVA was used for statistical comparison. Error bars represent the standard error of the mean.

We observed an expected and consistent pattern across all APL-MP groups: as grid resolution improved (i.e., as grid size decreased), detection sensitivity decreased and specificity increased (**Fig. 6**). For example, in the case of targeting both MUC1 and EPCAM antigens with (MUC1+EPCAM)-Protein L-MPs, the analysis revealed a mean detection sensitivity of 96% when utilizing the lowest grid resolution (10x10 grid or 178 µm resolution) and a mean detection sensitivity of 49% at the highest examined grid resolution (64x64 grid or 28 µm resolution) (**Fig. 6C**). These findings highlight the trade-off between sensitivity and specificity that occurs as the detection resolution becomes finer.

**Figure 6:**
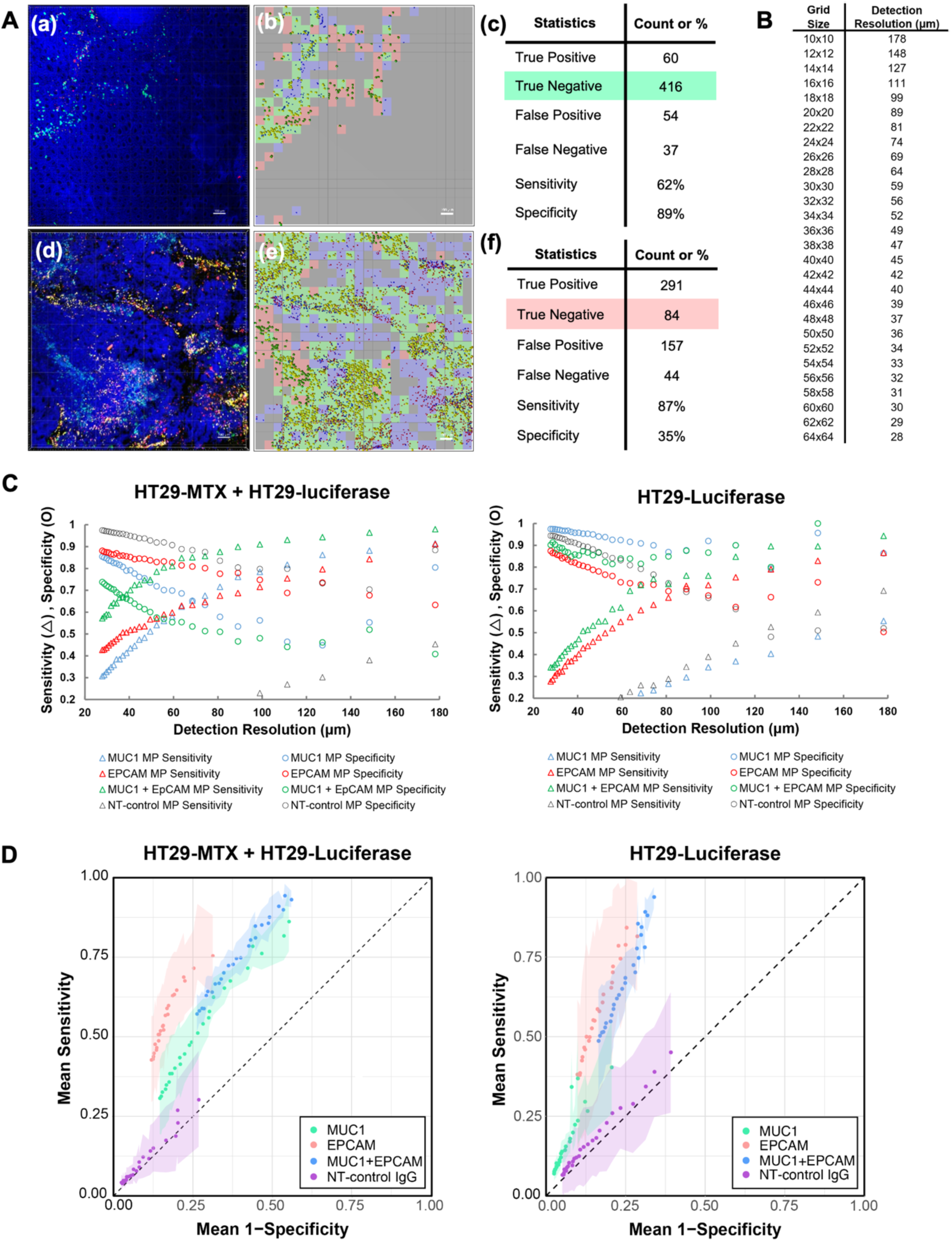
Quantification of sensitivity and specificity of APL-MPs detection using grid-based analysis. (**A**) Comparative image analysis of regions with (a,b) primarily normal tissue, and (d,e) primarily tumor-laden tissue. Confocal images in (a,d) show nuclei (DAPI, blue), HNA (green), EPCAM-APL-MPs (red), and MUC1-APL-MPs (yellow). Corresponding 24x24 grid maps, colored according to Fig. 5A(m), are shown in (b,e). Resultant statistics in (c,f) highlight higher specificity in majority normal tissue regions, and higher sensitivity in majority tumor-laden regions. (**B**) Table representing the relationship between grid size and detection resolution. (**C**) Sensitivity analysis: Comparison of individual and combined APL-MPs targeting MUC1^hi^/EPCAM^hi^ HT29-MTX/HT29-Luciferase HET-tumors (left) and MUC1^low^/EPCAM^hi^ HT29-Luciferase homogeneous tumors (right). (**D**) Receiver Operator Characteristic curves as a function of detection resolution for individual and combined APL-MPs targeting MUC1^hi^/EPCAM^hi^ HT29-MTX/HT29-Luciferase HET-tumors (left) and MUC1^low^/EPCAM^hi^ HT29-Luciferase homogeneous tumors (right).

Comparative analysis of detection sensitivity between individual APL-MPs and the combined (MUC1+EPCAM)-Protein L-MPs group demonstrated significantly higher sensitivities in the latter at certain detection resolutions. This enhancement in sensitivity was not observed consistently across all tested detection resolutions, highlighting the potential advantages of a combined approach utilizing both MUC1 and EPCAM targeting antibodies with Protein L-MPs, albeit with a dependence on specific detection resolution parameters (**Table SI.1**).

The increased sensitivity achieved when targeting two antigens concurrently in both MUC1^hi^/EPCAM^hi^ and MUC1^low^/EPCAM^hi^ tumors was accompanied by a decrease in specificity. Such a compromise is anticipated due to the inherent increases in signal and noise when using dual targeting agents. **Fig. 6C** shows the relationship between sensitivity and specificity at different detection resolutions.

## 4. Discussion

Visualization of tumors at the molecular level has applications in personalized treatment planning, surgical removal of lesions by identifying tumor margins, and early detection of occult lesions (6–9). Chromoendoscopy, in which topical dyes are applied to the colon surface, is an adjuvant endoscopic technique that can provide detailed contrast enhancement of dysplastic lesions (33). This technique has failed to gain acceptance as it did not offer an overall increase in the detection of colonic adenomas in a French cohort of patients (34,35). Fluorescence spectroscopy, in which the autofluorescence properties of tissues are analyzed, is an alternative dye-free imaging approach that can distinguish between normal and dysplastic tissues with high sensitivity (90%) and specificity (95%) (34), but unfortunately this analysis can only sample a small area of the colon (1-3 mm^2^). Furthermore, detection depends on placement of the probe at the right location and can only identify readily visible polypoid lesions (34). To improve detection and aid in delivery of theranostic agents, imaging approaches targeting molecules expressed on the surfaces of tumors are gaining traction.

Cancer is a continuously evolving and heterogeneous disease (1,36,37). Most colorectal tumors have high mutational heterogeneity at diagnosis, with thousands of mutations found in spatially separate regions of tumors (38). This genetic diversity has implications in cancer medicine, including in detection of CRC using targeted approaches. For this study, our first goal was to create CRC models that could be used to study the targeting of heterogeneous tumors, focusing on MUC1 and EPCAM. IF staining of CRC cell lines cultured in monolayer cultures revealed distinct patterns of antigen expression, consistent with their diverse tumor origins. These findings highlight the heterogeneity in antigen expression within CRC cell lines and lay the foundation for subsequent *in vitro* and *in vivo* experiments using HET-tumoroids. The irregular expression pattern of EPCAM in HT29-MTX tumoroids may be attributed to the size and presence of MUC1 protein, which could interfere with detection of the EPCAM antigen. MUC1 is a remarkably large and heavily glycosylated mucin glycoprotein with a glycosylated peptide backbone that extends approximately 200-500 nanometers above the cell surface (14). Carcinoembryonic antigen (CEA) is another widely used marker for CRC and could have been added to MUC1 and EPCAM to create a trivalent detection system (13). However, although CEA was detected by IHC in the tissue specimens (**Fig. SI.1.A.**), we did not find sufficient and reliable CEA expression across multiple CRC cell lines ((**Fig. SI.1.B.**) and therefore chose not to develop CEA-functional APL-MPs.

We tagged MPs with fluorophores either indirectly using secondary antibodies or directly using click chemistry to attach Alexa Fluor™ fluorophores to antibodies. The click chemistry labeling approach offers several advantages for labeling IgG antibodies and is often used to label antibody-drug conjugates (ADCs) (39). First, it enables the direct conjugation of the fluorophore to the Fc region of the primary antibody, thus securing the fluorophore away from the Fab region, where Protein L binds, and where the antibody recognizes antigen. Second, the click chemistry labeling method eliminates the need for a secondary antibody, which can lead to non-specific binding and increased background signal. This is particularly important in experiments that require the use of more than one antibody made in the same host tissue, as it reduces the likelihood of non-specific binding of secondary and primary antibodies. This simple, rapid, and versatile labeling approach provides a broad range of fluorescent dyes to choose from, allowing for a more comprehensive visualization and targeting of heterogeneity within tumors. By using a mixture of MPs labeled with different antibodies and fluorophores of different colors, we were able to capture the molecular signatures of two different antigens simultaneously.

We determined that orthotopic human CRC tumors display MUC1 and/or EPCAM, as appropriate to the injected source cells, on the luminal tumor surface in a manner that was accessible to the APL-MPs. Notably, the resultant tumors did not adopt a pedunculated morphology, but were comparably flat across the luminal surface, gradually expanding into the open space. This has further value in replicating both heterogeneous biomarker expression and the uncommon tumor morphology that is more difficult to observe by standard colonoscopy.

To evaluate the sensitivity and specificity of detection, we employed a grid-based analysis approach to examine the positional relationships between segmented objects within a grid box. This analysis considered the localization of particles in general proximity (same grid-box) as human cancer nuclei to quantitatively assess the binding sensitivity and specificity of the particles. The results consistently demonstrated a noticeable decrease in detection sensitivity and an increase in specificity as the grid resolution became finer. Comparative analysis of detection sensitivity between individual APL-MPs (MUC1 or EPCAM, or IgG control alone) and the combined (MUC1+EPCAM)-APL-MPs group revealed higher sensitivities at certain detection resolutions. Interestingly, as the grid sizes decreased and the detection resolutions increased, the sensitivities began to converge. These results imply that the choice of grid size and detection resolution is crucial for determining the sensitivity of detection for each antigen.

APL-MPs have promising applications in various areas including surgical margin detection, personalized treatment planning, and theranostic applications. By specifically targeting tumor-associated antigens, APL-MPs potentially can aid in accurate tumor detection and margin assessment during surgical procedures. This concept has been explored in previous studies using magnetic particle imaging or spectroscopic analysis of tissue autofluorescence for identification of residual tumors in breast-conserving surgery (40,41). Additionally, the ability of APL-MPs to recognize specific antigens opens avenues for personalized therapy, where targeted treatments can be tailored based on an individual’s tumor profile.

Another potential application of APL-MPs is the detection of heterogeneous lesions using various molecular imaging techniques, such as hyperpolarized (HP) MRI, confocal laser endomicroscopy (CLE), and Raman spectroscopy. One study demonstrated the feasibility of detecting MUC1 expressing tumors *in vivo* using antibody-functionalized HP silicon particles in a CRC mouse model (42)(43). In this study, 2 µm HP silicon particles functionalized with an antibody against MUC1 were administered to human-MUC1 expressing mice via the rectum, and the particles were detected using MRI at the tumor site. HP-MRI offers enhanced sensitivity and signal-to-noise (SNR) ratio by enhancing the nuclear spin alignment of the MR detectable MPs. Hyperpolarization of silicon requires extended exposure of the agent to near-zero Kelvin temperature. In parallel to Figure SI.2., we verified (not shown) that Protein L binding of control IgG1 remained unaffected after such temperature exposure, suggesting the feasibility in future MRI studies of APL-MP analogs based on HP silicon.

In addition to HP-MRI, CLE is another emerging technology that enables real-time histological assessment of the mucosal layer of the gastrointestinal (GI) tract. CLE utilizes fluorescent probes to selectively highlight specific tissue features, allowing for detailed visualization of the tumor. In a separate study, CLE was used to detect fluorescein-conjugated peptides administered topically to dysplastic colon, with a sensitivity of 81% and specificity of 82% (44). The Protein L labeling approach also can be extended to Raman scattering (SERS) MPs, allowing conjugation with antibodies to a diverse panel of tumor-specific antigens. These particles then can be visualized with a Raman endoscope inserted through the accessory channel of a conventional endoscope (45). The flexibility of Protein L as a surface adapter allows for an array of particle cargos and mono- or multivalent surface functionalization methods to be used for any cancer signature.

## 5. Conclusions

In summary, our study contributes to a deeper understanding of antigen expression heterogeneity in CRC and highlights the potential of targeted detection using APL-MPs. Our findings highlight the significance of considering tumor heterogeneity and utilizing multi-antigen targeting strategies to enhance detection sensitivity in studies involving antibody-based imaging or therapy. The combination of MUC1-specific and EPCAM-specific targeting particles broadened the identification of diverse model tumors and demonstrated their potential value in clinical samples for which antigen expression is unknown and/or dynamic. The flexibility of the surface coupling concept enables its application to an even broader array of antigens and can translate to other microparticle compositions as needed. Overall, these findings pave the way for improved antigen-targeted diagnostics and personalized treatment strategies for patients with CRC.

## Supplementary Materials

Supplementary data for this article are available at Cancer Research Online

## Author Contributions

**S. Ramezani:** Conceptualization, investigation, data curation, formal analysis, methodology, visualization, writing–original draft, writing–review, and editing. **N. M. Zacharias:** Investigation, methodology, visualization, writing, review, and editing. **W. Norton:** Investigation, methodology, writing–review and editing, **J. S. Davis:** Methodology, writing–review and editing. **A. Dominic:** Data curation, formal analysis, writing–original draft, writing–review, and editing. **R. Armijo:** investigation and methodology. **M. Wang:** investigation and methodology. **R. E. Wendt:** Formal analysis, writing the original draft. **D. D. Carson:** Conceptualization, formal analysis, methodology, writing, reviewing, and editing. **D. A. Harrington:** conceptualization, data curation, formal analysis, methodology, visualization, writing–original draft, writing–review and editing, resources, funding acquisition, project administration **M. C. Farach-Carson:** conceptualization, data curation, formal analysis, methodology, visualization, writing–original draft, writing–review and editing, resources, funding acquisition, project administration **P. K. Bhattacharya:** conceptualization, formal analysis, methodology, writing–review and editing, resources, funding acquisition, project administration

## Institutional Review Board Statement

Not applicable.

## Informed Consent Statement

Not applicable.

## Acknowledgments

We gratefully acknowledge Dr. Lissette Cruz for assistance in developing the tumoroids methodology, Dr. Marty Pagel for valuable conversations on data analysis, and Dr. Zhiqiang Ahn for suggesting the use of protein L. We also gratefully acknowledge the team contributions of past members of the Bhattacharya, Carson, Farach-Carson, Harrington, and Wu laboratories. Confocal fluorescence microscopy, spectroscopy for ELISA, and image analysis workstations were used as shared instrumentation at the Center for Craniofacial Research Instrumentation Core, School of Dentistry, UTHealth Houston. This work was supported by the following financial support: S.R. acknowledges the UTHealth Innovation for Cancer Prevention Research Training Program (Cancer Prevention and Research Institute of Texas grant #RP160015). P.B. acknowledges the Cancer Prevention and Research Institute of Texas (CPRIT; RP220270), US National Institute of Biomedical Imaging and Bioengineering (NIBIB, R21EB031217), US Department of Defense (W81XWH-21-1-0763, HT9425310664), Institutional Research Grants, and a Startup grant from MD Anderson Cancer Center. D.A.H. acknowledges startup funds from The University of Texas Health Science Center at Houston. M.C.F.C acknowledges a Translational Faculty Science and Technology Acquisition and Retention (STARs) Program Award of the University of Texas System and Institutional Start-up Funds. Disclaimer: The content is solely the responsibility of the authors and does not necessarily represent the official views of the Cancer Prevention and Research Institute of Texas or the National Institutes of Health.

## References

1. Burrell RA, McGranahan N, Bartek J, Swanton C. The causes and consequences of genetic heterogeneity in cancer evolution. Nature. Nature Publishing Group; 2013;501:338–45.

2. Dagogo-Jack I, Shaw AT. Tumour heterogeneity and resistance to cancer therapies. Nat Rev Clin Oncol. 2018;15:81–94.

3. Zahir N, Sun R, Gallahan D, Gatenby RA, Curtis C. Characterizing the ecological and evolutionary dynamics of cancer. Nat Genet. 2020;52:759–67.

4. McGranahan N, Swanton C. Clonal Heterogeneity and Tumor Evolution: Past, Present, and the Future. Cell. 2017;168:613–28.

5. Zhou K, Li G, Pan R, Xin S, Wen W, Wang H, et al. Preclinical evaluation of AGTR1-Targeting molecular probe for colorectal cancer imaging in orthotopic and liver metastasis mouse models. Eur J Med Chem. 2024;271:116452.

6. Schaafsma BE, Mieog JSD, Hutteman M, van der Vorst JR, Kuppen PJK, Löwik CWGM, et al. The clinical use of indocyanine green as a near-infrared fluorescent contrast agent for image-guided oncologic surgery. J Surg Oncol. 2011;104:323–32.

7. Troyan SL, Kianzad V, Gibbs-Strauss SL, Gioux S, Matsui A, Oketokoun R, et al. The FLARE^TM^ Intraoperative Near-Infrared Fluorescence Imaging System: A First-in-Human Clinical Trial in Breast Cancer Sentinel Lymph Node Mapping. Ann Surg Oncol. 2009;16:2943–52.

8. Bhuvaneswari R, Thong PS-P, Gan-Yap YY, Soo K-C, Olivo MC. Evaluation of hypericin-mediated photodynamic therapy in combination with angiogenesis inhibitor bevacizumab using in vivo fluorescence confocal endomicroscopy. J Biomed Opt. SPIE; 2010;15:011114.

9. Khemthongcharoen N, Jolivot R, Rattanavarin S, Piyawattanametha W. Advances in imaging probes and optical microendoscopic imaging techniques for early in vivo cancer assessment. Adv Drug Deliv Rev. 2014;74:53–74.

10. Tainsky MA. Genomic and proteomic biomarkers for cancer: a multitude of opportunities. Biochim Biophys Acta. 2009;1796:176–93.

11. Pal M, Muinao T, Boruah HPD, Mahindroo N. Current advances in prognostic and diagnostic biomarkers for solid cancers: Detection techniques and future challenges. Biomed Pharmacother. 2022;146:112488.

12. Nilson BH, Solomon A, Björck L, Akerström B. Protein L from Peptostreptococcus magnus binds to the kappa light chain variable domain. J Biol Chem. 1992;267:2234–9.

13. Ramezani S, Parkhideh A, Bhattacharya P, Farach-Carson MC (Cindy), Harrington DA. Beyond Colonoscopy: Exploring New Cell Surface Biomarkers for Detection of Early, Heterogenous Colorectal Lesions. Front Oncol [Internet]. Frontiers; 2021 [cited 2021 Apr 26];11. Available from: https://www.frontiersin.org/articles/10.3389/fonc.2021.657701/abstract

14. Brayman M, Thathiah A, Carson DD. MUC1: A multifunctional cell surface component of reproductive tissue epithelia. Reprod Biol Endocrinol RBE. 2004;2.

15. Cascio S, Finn OJ. Complex of MUC1, CIN85 and Cbl in Colon Cancer Progression and Metastasis. Cancers. 2015;7:342–52.

16. Holst S, Wuhrer M, Rombouts Y. Glycosylation characteristics of colorectal cancer. Adv Cancer Res. 2015;126:203–56.

17. Pudakalakatti S, Enriquez J, McGowan C, Ramezani S, Davis J, Zacharias N, et al. Hyperpolarized MRI with Silicon Micro and Nanoparticles: Principles and Applications. WIREs Nanomedicine Nanotechnolgoy. 2021;

18. Wang H-S, Wang L-H. The expression and significance of Gal-3 and MUC1 in colorectal cancer and colon cancer. OncoTargets Ther. 2015;8:1893–8.

19. Betge J, Schneider NI, Harbaum L, Pollheimer MJ, Lindtner RA, Kornprat P, et al. MUC1, MUC2, MUC5AC, and MUC6 in colorectal cancer: expression profiles and clinical significance. Virchows Arch Int J Pathol. 2016;469.

20. Maetzel D, Denzel S, Mack B, Canis M, Went P, Benk M, et al. Nuclear signalling by tumour-associated antigen EpCAM. Nat Cell Biol. 2009;11:162–71.

21. Trzpis M, McLaughlin PMJ, Leij LMFH de, Harmsen MC. Epithelial Cell Adhesion Molecule: More than a Carcinoma Marker and Adhesion Molecule. Am J Pathol. Elsevier; 2007;171:386–95.

22. van der Gun BTF, Melchers LJ, Ruiters MHJ, de Leij LFMH, McLaughlin PMJ, Rots MG. EpCAM in carcinogenesis: the good, the bad or the ugly. Carcinogenesis. 2010;31:1913–21.

23. Fong D, Moser P, Kasal A, Seeber A, Gastl G, Martowicz A, et al. Loss of membranous expression of the intracellular domain of EpCAM is a frequent event and predicts poor survival in patients with pancreatic cancer. Histopathology. John Wiley & Sons, Ltd; 2014;64:683–92.

24. Gastl G, Spizzo G, Obrist P, Dünser M, Mikuz G. Ep-CAM overexpression in breast cancer as a predictor of survival. Lancet Lond Engl. 2000;356:1981–2.

25. Spizzo G, Gastl G, Obrist P, Went P, Dirnhofer S, Bischoff S, et al. High Ep-CAM Expression is Associated with Poor Prognosis in Node-positive Breast Cancer. Breast Cancer Res Treat. 2004;86:207–13.

26. Seeber A, Untergasser G, Spizzo G, Terracciano L, Lugli A, Kasal A, et al. Predominant expression of truncated EpCAM is associated with a more aggressive phenotype and predicts poor overall survival in colorectal cancer. Int J Cancer. 2016;139:657–63.

27. Massoner P, Thomm T, Mack B, Untergasser G, Martowicz A, Bobowski K, et al. EpCAM is overexpressed in local and metastatic prostate cancer, suppressed by chemotherapy and modulated by MET-associated miRNA-200c/205. Br J Cancer. 2014;111:955–64.

28. Spizzo G, Went P, Dirnhofer S, Obrist P, Moch H, Baeuerle PA, et al. Overexpression of epithelial cell adhesion molecule (Ep-CAM) is an independent prognostic marker for reduced survival of patients with epithelial ovarian cancer. Gynecol Oncol. 2006;103:483–8.

29. Fong D, Steurer M, Obrist P, Barbieri V, Margreiter R, Amberger A, et al. Ep-CAM expression in pancreatic and ampullary carcinomas: frequency and prognostic relevance. J Clin Pathol. 2008;61:31–5.

30. Sam-Yellowe TY. Immunology: Overview and Laboratory Manual [Internet]. Cham: Springer International Publishing; 2021 [cited 2025 Jul 28]. Available from: https://link.springer.com/10.1007/978-3-030-64686-8

31. Bettenworth D, Mücke MM, Schwegmann K, Faust A, Poremba C, Schäfers M, et al. Endoscopy-guided orthotopic implantation of colorectal cancer cells results in metastatic colorectal cancer in mice. Clin Exp Metastasis. 2016;33:551–62.

32. De Château M, Nilson BHK, Erntell M, Myhre E, Magnusson CGM, Åkerström B, et al. On the Interaction between Protein L and Immunoglobulins of Various Mammalian Species. Scand J Immunol. 1993;37:399–405.

33. Buchner AM. The Role of Chromoendoscopy in Evaluating Colorectal Dysplasia. Gastroenterol Hepatol. 2017;13:336–47.

34. DaCosta RS, Wilson BC, Marcon NE. Optical techniques for the endoscopic detection of dysplastic colonic lesions. Curr Opin Gastroenterol. 2005;21:70–9.

35. Le Rhun M, Coron E, Parlier D, Nguyen J, Canard J, Alamdari A, et al. High Resolution Colonoscopy With Chromoscopy Versus Standard Colonoscopy for the Detection of Colonic Neoplasia: A Randomized Study. Clin Gastroenterol Hepatol. 2006;4:349–54.

36. Vendramin R, Litchfield K, Swanton C. Cancer evolution: Darwin and beyond. EMBO J. John Wiley & Sons, Ltd; 2021;40:e108389.

37. Merlo LMF, Pepper JW, Reid BJ, Maley CC. Cancer as an evolutionary and ecological process. Nat Rev Cancer. Nature Publishing Group; 2006;6:924–35.

38. Uchi R, Takahashi Y, Niida A, Shimamura T, Hirata H, Sugimachi K, et al. Integrated Multiregional Analysis Proposing a New Model of Colorectal Cancer Evolution. Gerlinger M, editor. PLOS Genet. 2016;12:e1005778.

39. Dudchak R, Podolak M, Holota S, Szewczyk-Roszczenko O, Roszczenko P, Bielawska A, et al. Click chemistry in the synthesis of antibody-drug conjugates. Bioorganic Chem. 2024;143:106982.

40. Shipp DW, Rakha EA, Koloydenko AA, Macmillan RD, Ellis IO, Notingher I. Intra-operative spectroscopic assessment of surgical margins during breast conserving surgery. Breast Cancer Res. 2018;20:69.

41. Mason EE, Mattingly E, Herb K, Śliwiak M, Franconi S, Cooley CZ, et al. Concept for using magnetic particle imaging for intraoperative margin analysis in breast-conserving surgery. Sci Rep. 2021;11:13456.

42. In Vivo Molecular Imaging of MUC1-Expressing Colorectal Tumors Using Targeted Hyperpolarized Silicon Particles. Jt Annu Meet ISMRM-ESMRMB [Internet]. Paris, France; 2018. page 0105. Available from: https://cds.ismrm.org/protected/18MProceedings/PDFfiles/0105.html

43. McCowan CV, Salmon D, Hu J, Pudakalakatti S, Whiting N, Davis JS, et al. Post-Acquisition Hyperpolarized 29Silicon Magnetic Resonance Image Processing for Visualization of Colorectal Lesions Using a User-Friendly Graphical Interface. Diagn Basel Switz. 2022;12:610.

44. Hsiung P, Hardy J, Friedland S, Soetikno R, Du CB, Wu APW, et al. Detection of colonic dysplasia in vivo using a targeted fluorescent septapeptide and confocal microendoscopy. Nat Med. 2008;14:454–8.

45. Zavaleta CL, Garai E, Liu JTC, Sensarn S, Mandella MJ, Sompel de DV, et al. A Raman-based endoscopic strategy for multiplexed molecular imaging. Proc Natl Acad Sci. 2013;110:2288–97.

