## Supplemental Information for "Antibody-Protein L Functionalized Microparticles for Detection of Surface Markers in Heterogeneous Colorectal Lesions"

### Supplementary Information

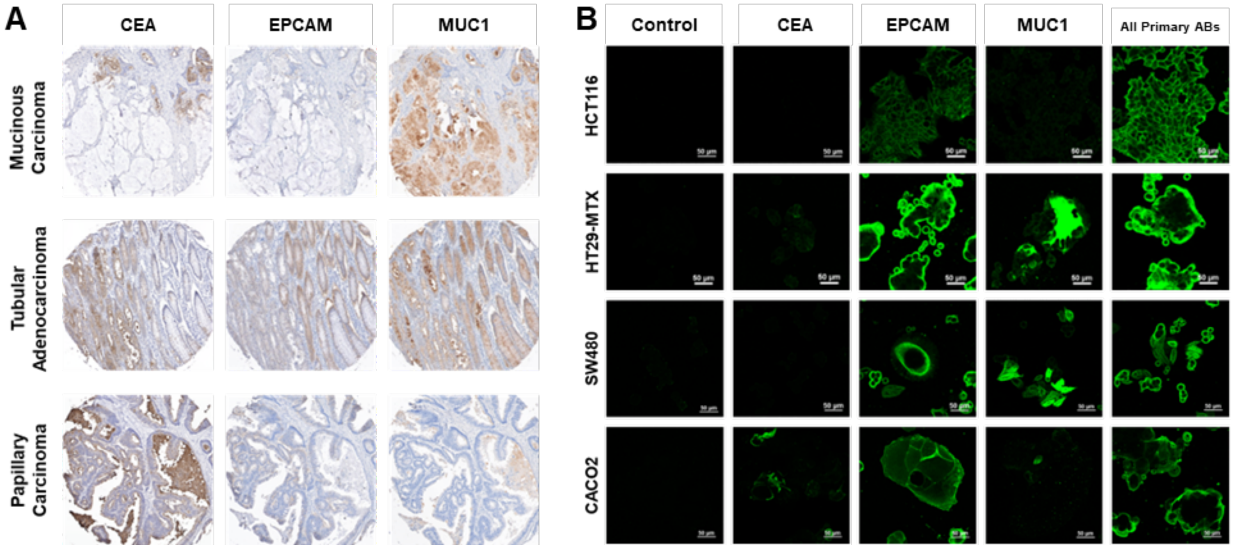


**Figure SI.1. Immunohistochemistry (IHC) and immunofluorescence (IF) staining show that surface markers MUC1, EPCAM, and CEA are heterogeneously expressed.** (**A**) IHC staining (brown = positive) for CEA, EPCAM and MUC1 in mucinous adenocarcinoma, tubular adenocarcinoma, and papillary carcinoma human tissue samples. (**B**) IF staining (green = positive) for CEA, EPCAM, and MUC1 in CRC cell lines, revealing heterogeneous expression of surface antigens in four CACO2, SW480, HT29-MTX, and HCT116 cell lines.

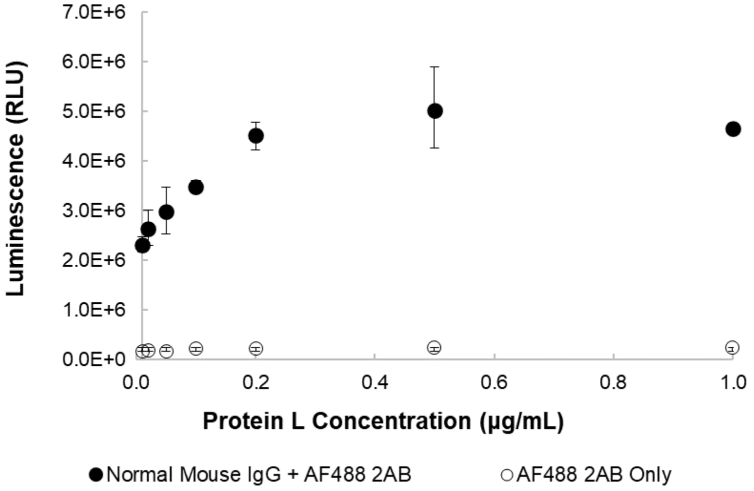


**Figure SI.2. Assessment of the binding capacity of Protein L to non-targeted (NT) mouse IgG1 control antibody using an ELISA assay.** The saturation point for Protein L binding was observed at a concentration of 0.2 μg/mL. The error bars are standard deviations of three separate measurements.

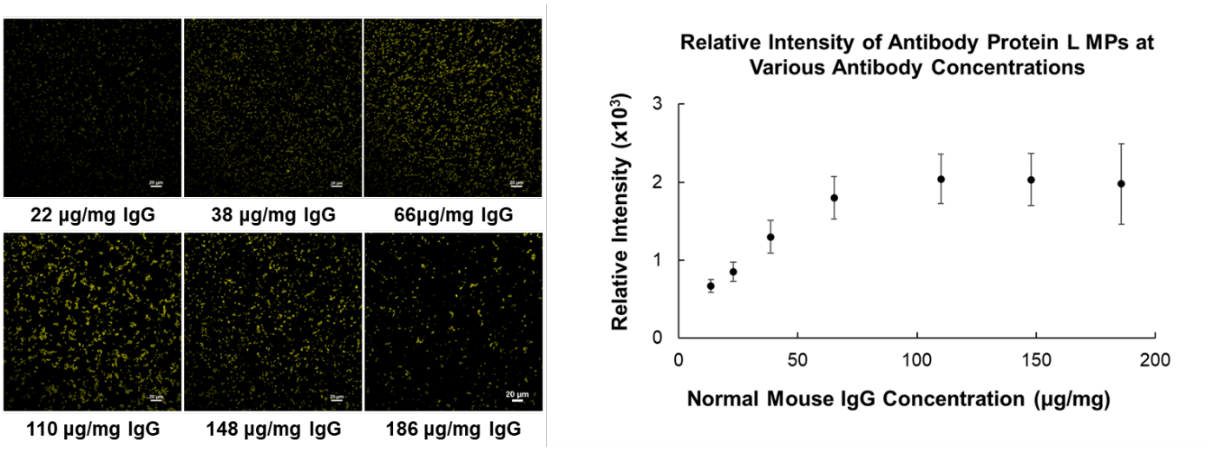


**Figure SI.3. Saturation of Fluorescence Signal in MPs with Increasing Antibody Loading.** Sequential loading of Alexa Fluor™ 647 labeled NT-control mouse IgG1 antibody onto Protein L MPs. The left panel demonstrates confocal microscope images of the MPs as the loading of antibody increases, showing clear saturation of the signal around 110 μg of IgG1 antibody per mg of MP. The right panel provides a graphical representation of the fluorescence signal intensity in relation to the antibody concentration, confirming the observed saturation point. The error bars are standard deviation of ten random measurements of fluorescent intensity in each image.

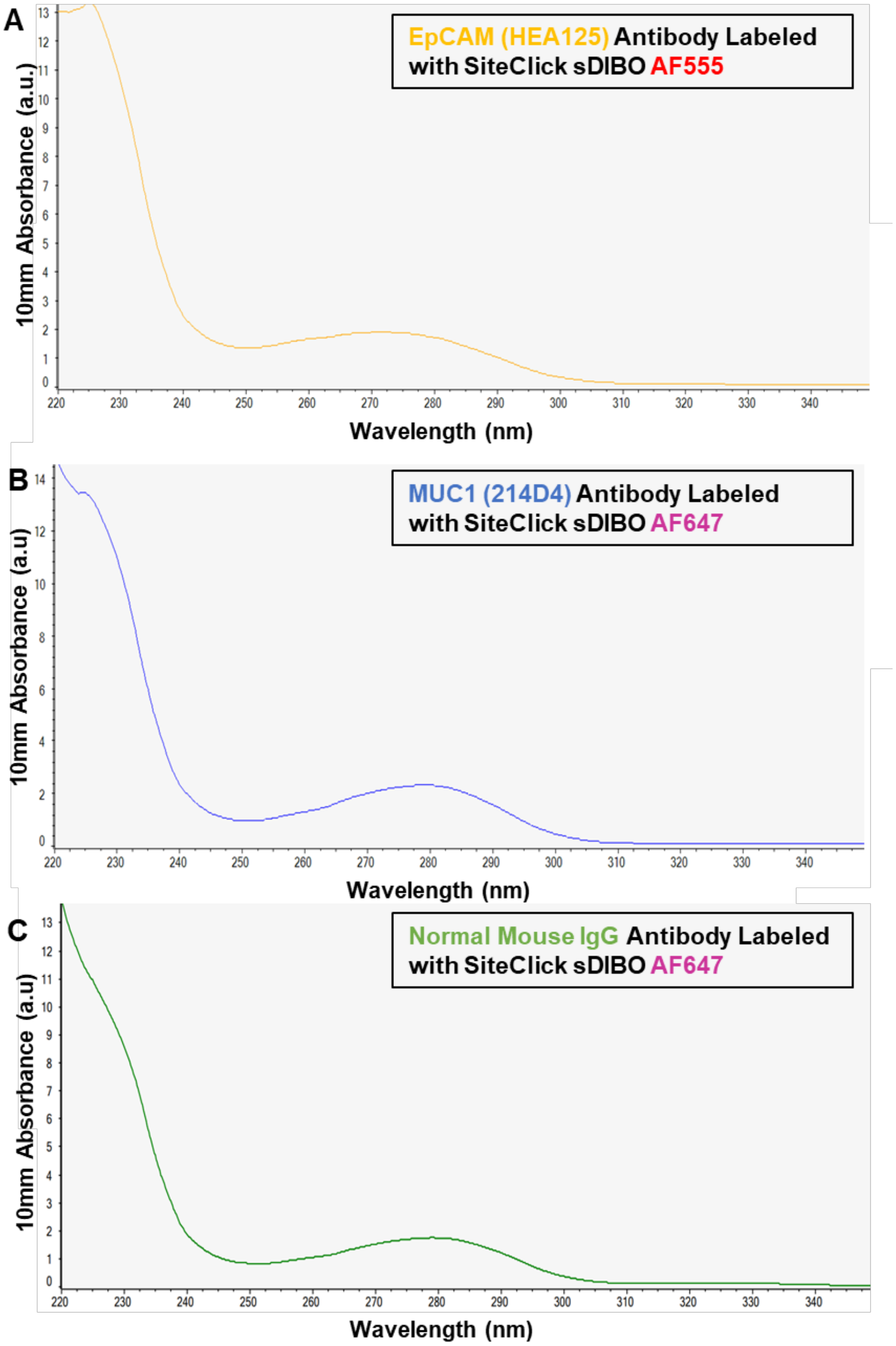


**Figure SI.4. Absorbance spectra of antibodies labeled with Alexa Fluor™ fluorophores, as measured by the NanoDrop spectrophotometer.** (A) The absorbance spectrum of EpCAM (HEA125) antibody labeled with Alexa Fluor™ 555, (B) MUC1 (214D4) antibody labeled with Alexa Fluor™ 647, and (C) NT-control mouse IgG antibody labeled with Alexa Fluor™ 647.

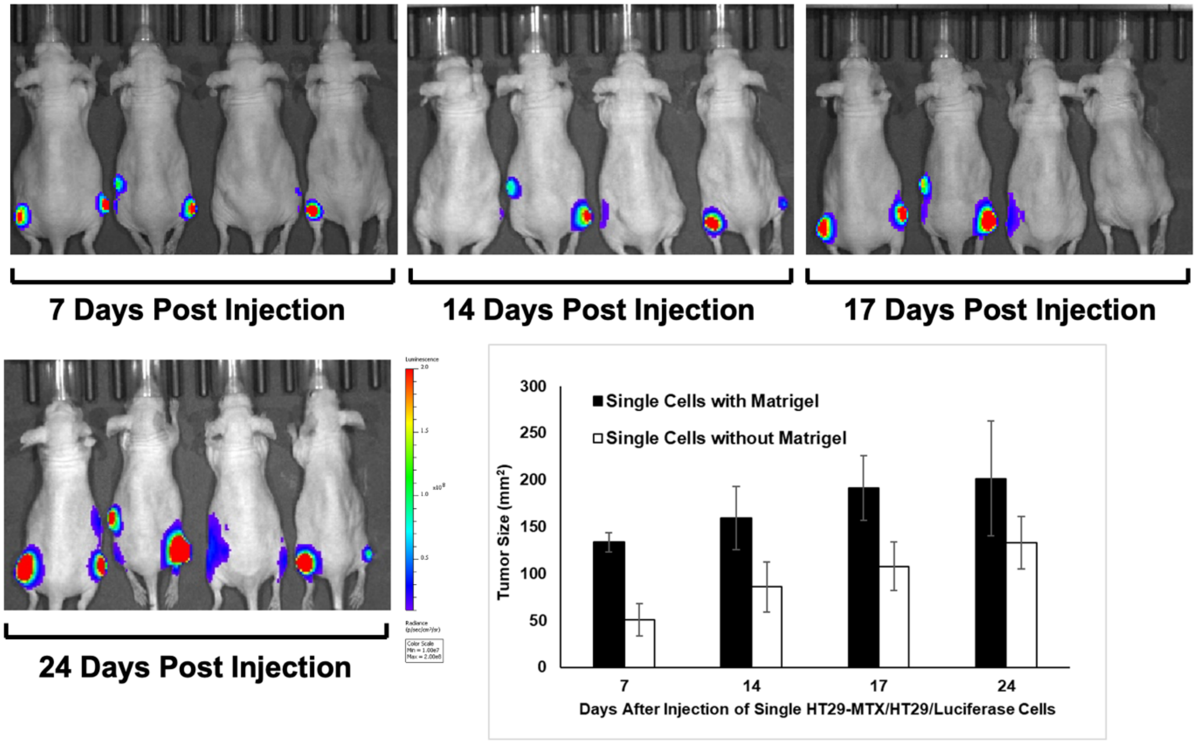


**Figure SI.5. 9 weeks-old mice injected with HT29-MTX and HT29-Luciferase CRC single cells.** A mixture of HT29-MTX (70%)/HT29-Luciferase (30%) cells in 1:1 PBS/Matrigel were injected into the left flank and cells in PBS alone (without Matrigel) were injected into the right flank of nude mice. Bioluminescence images of mice 10 minutes post-luciferin injection are shown after 7, 14, 17, and 24 days after implantation of cells. Tumor sizes determined by calipers at the same time points are illustrated in the bar graph.

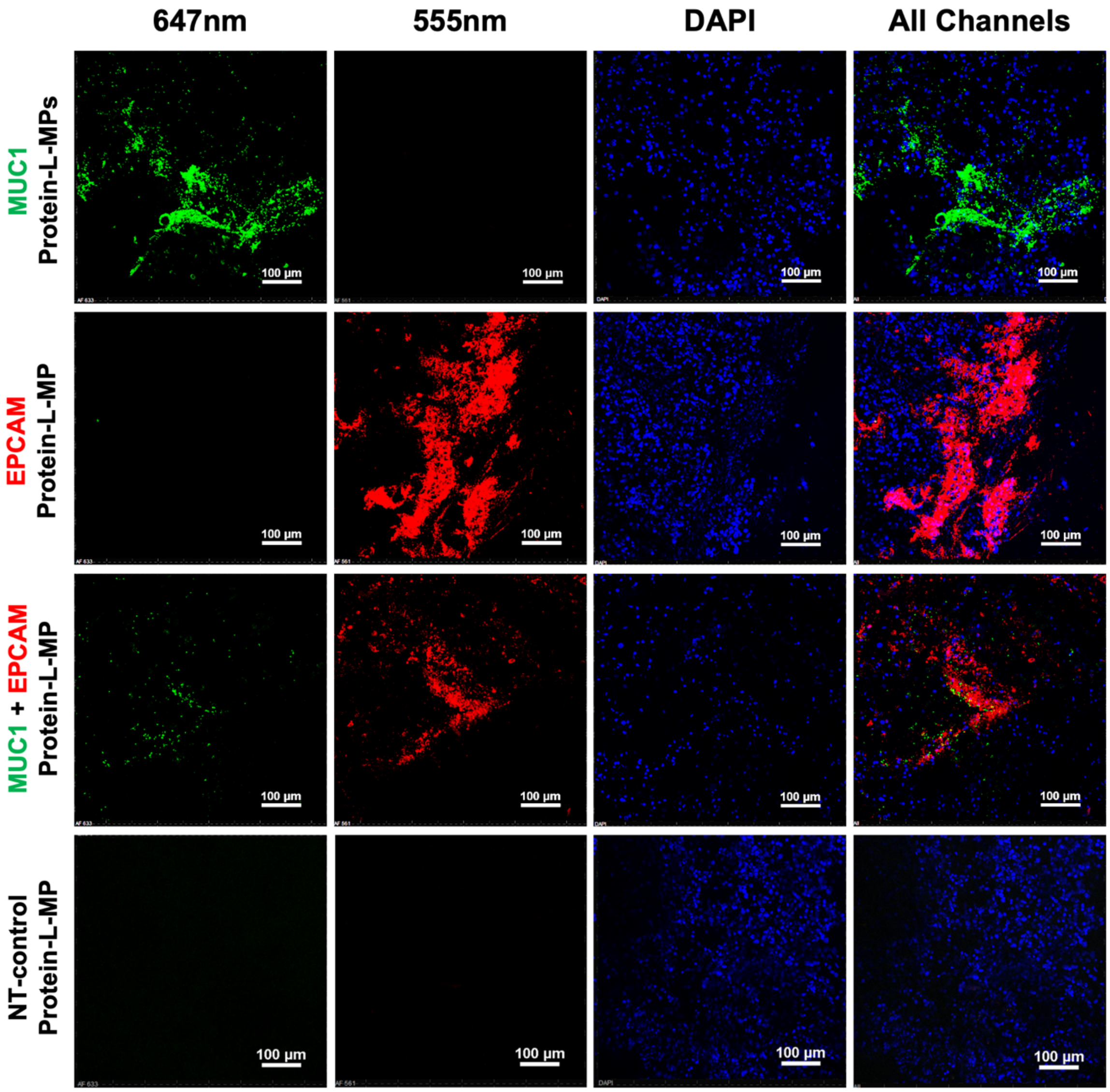


**Figure SI.6. Antibody-Protein L-MPs recognize human antigens on flank tumors.** Tissue sections from flank HET-tumors were incubated with MPs targeting either MUC1, EPCAM, a combination of both MPs, and control IgG. Specific binding of the MPs to the HET-tumoroids was observed, with non-overlapping signals in the overlaid image. Negligible signal was observed for the NT-control IgG antibody-Protein L MPs in the flank tumors.

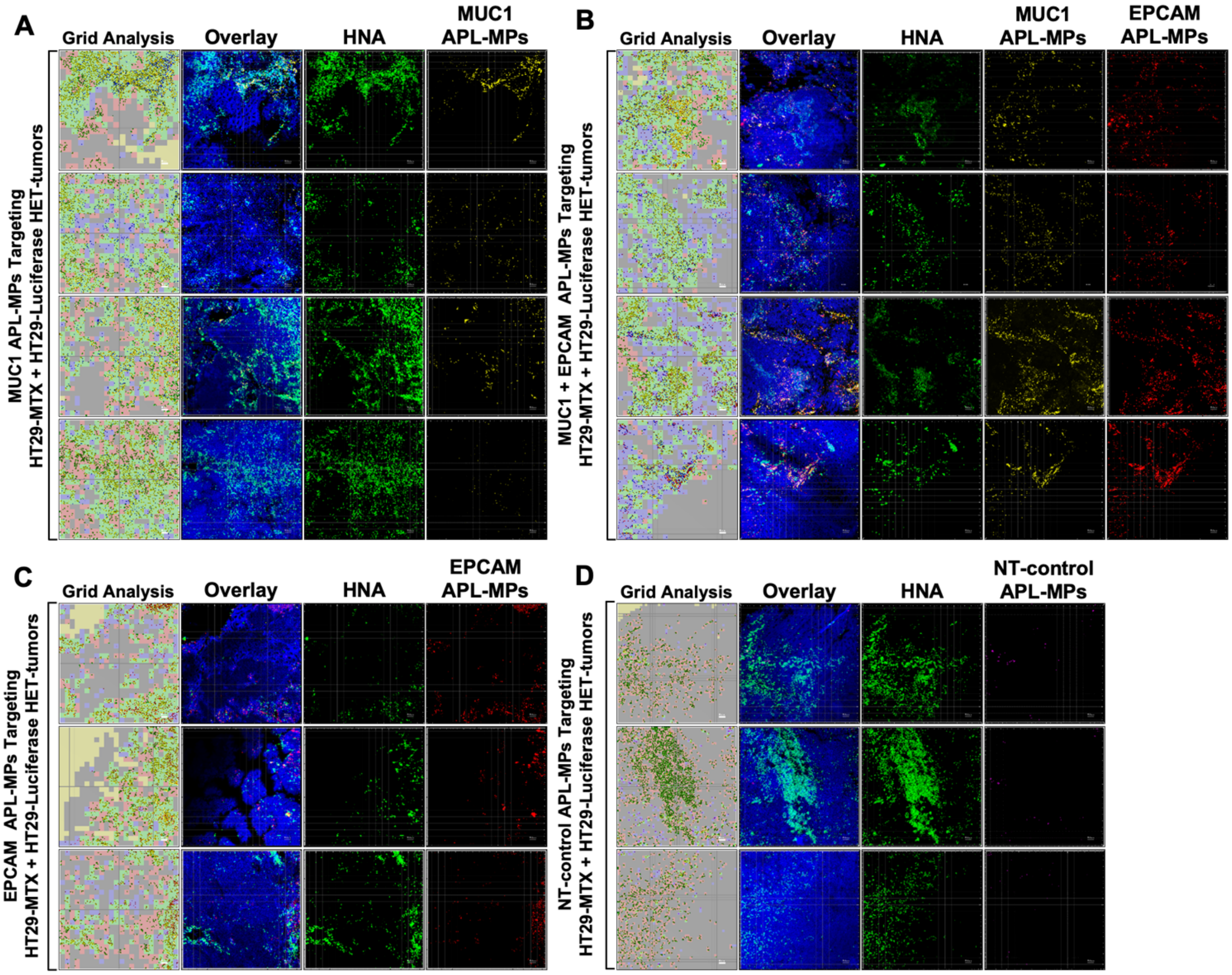


**Figure SI.7. Colonic tissues incubated with APL-MPs *in vivo* demonstrate specific antigen targeting.** Confocal images of HT29-MTX + HT29-Luciferase HET-tumors being detected with (**A**) MUC1, (**B**) MUC1 + EPCAM, and (**C**) EPCAM APL-MPs simultaneously. MUC1 APL-MPs showed specific localization in regions of MUC1$^{hi}$ HT29-MTX-induced homogeneous tumors, while EPCAM APL-MPs were densely concentrated near EPCAM$^{hi}$ HT29-Luciferase-induced homogeneous tumors, highlighting their targeting capabilities. (**D**) In contrast, NT-control IgG1 APL-MPs exhibited minimal to no binding to CRC cells, indicating their lack of specific interaction.

HT29-MTX/HT29-Luciferase
MUC1[hi]/EPCAM[hi] HET-tumors
Adjusted P values for sensitivity at various grid sizes

| Grid Size | MUC1 + EPCAM vs. MUC1 | MUC1 + EPCAM vs. EPCAM | MUC1 + EPCAM vs. NT-control | MUC1 vs. EPCAM | MUC1 vs. NT-control | EPCAM vs. NT-control |
|---|---|---|---|---|---|---|
| 8x8 | 0.9361 (ns) | 0.962 (ns) | 0.0029 (**) | >0.9999 (ns) | 0.0064 (**) | 0.0091 (**) |
| 10x10 | 0.8297 (ns) | 0.8434 (ns) | 0.0004 (***) | >0.9999 (ns) | 0.0011 (**) | 0.002 (**) |
| 12x12 | 0.7126 (ns) | 0.4939 (ns) | 0.0002 (***) | 0.9647 (ns) | 0.0006 (***) | 0.0019 (**) |
| 14x14 | 0.6312 (ns) | 0.2372 (ns) | <0.0001 (****) | 0.7985 (ns) | <0.0001 (****) | 0.0004 (***) |
| 16x16 | 0.3211 (ns) | 0.1013 (ns) | <0.0001 (****) | 0.7903 (ns) | <0.0001 (****) | 0.0003 (***) |
| 18x18 | 0.2073 (ns) | 0.05 (*) | <0.0001 (****) | 0.7068 (ns) | <0.0001 (****) | 0.0002 (***) |
| 20x20 | 0.1191 (ns) | 0.022 (*) | <0.0001 (****) | 0.6088 (ns) | <0.0001 (****) | <0.0001 (****) |
| 22x22 | 0.0413 (*) | 0.0186 (*) | <0.0001 (****) | 0.8794 (ns) | <0.0001 (****) | <0.0001 (****) |
| 24x24 | 0.0154 (*) | 0.0099 (**) | <0.0001 (****) | 0.941 (ns) | <0.0001 (****) | <0.0001 (****) |
| 26x26 | 0.0128 (*) | 0.0114 (*) | <0.0001 (****) | 0.9823 (ns) | <0.0001 (****) | <0.0001 (****) |
| 28x28 | 0.0044 (**) | 0.0085 (**) | <0.0001 (****) | 0.9997 (ns) | <0.0001 (****) | <0.0001 (****) |
| 30x30 | 0.0037 (**) | 0.0106 (*) | <0.0001 (****) | 0.9846 (ns) | <0.0001 (****) | <0.0001 (****) |
| 32x32 | 0.0029 (**) | 0.0115 (*) | <0.0001 (****) | 0.944 (ns) | <0.0001 (****) | <0.0001 (****) |
| 34x34 | 0.0059 (**) | 0.0212 (*) | <0.0001 (****) | 0.9573 (ns) | <0.0001 (****) | <0.0001 (****) |
| 36x36 | 0.0066 (**) | 0.0482 (*) | <0.0001 (****) | 0.7839 (ns) | 0.0001 (***) | <0.0001 (****) |
| 38x38 | 0.0037 (**) | 0.0243 (*) | <0.0001 (****) | 0.8181 (ns) | 0.0003 (***) | 0.0002 (***) |
| 40x40 | 0.0046 (**) | 0.0362 (*) | <0.0001 (****) | 0.7568 (ns) | 0.0002 (***) | 0.0001 (***) |
| 42x42 | 0.0033 (**) | 0.0373 (*) | <0.0001 (****) | 0.6346 (ns) | 0.0005 (***) | 0.0002 (***) |
| 44x44 | 0.005 (**) | 0.067 (ns) | <0.0001 (****) | 0.5691 (ns) | 0.0007 (***) | 0.0002 (***) |
| 46x46 | 0.0056 (**) | 0.0907 (ns) | <0.0001 (****) | 0.4993 (ns) | 0.0013 (**) | 0.0004 (***) |
| 48x48 | 0.0046 (**) | 0.1092 (ns) | <0.0001 (****) | 0.3796 (ns) | 0.0014 (**) | 0.0003 (***) |
| 50x50 | 0.006 (**) | 0.1039 (ns) | <0.0001 (****) | 0.4762 (ns) | 0.0032 (**) | 0.0007 (***) |
| 52x52 | 0.0097 (**) | 0.134 (ns) | <0.0001 (****) | 0.5465 (ns) | 0.0027 (**) | 0.0007 (***) |
| 54x54 | 0.0073 (**) | 0.1272 (ns) | <0.0001 (****) | 0.472 (ns) | 0.0066 (**) | 0.0013 (**) |
| 56x56 | 0.0072 (**) | 0.1349 (ns) | <0.0001 (****) | 0.445 (ns) | 0.0067 (**) | 0.0012 (**) |
| 58x58 | 0.0098 (**) | 0.2176 (ns) | <0.0001 (****) | 0.3776 (ns) | 0.0072 (**) | 0.0011 (**) |
| 60x60 | 0.008 (**) | 0.1737 (ns) | <0.0001 (****) | 0.3931 (ns) | 0.0081 (**) | 0.0013 (**) |
| 62x62 | 0.0083 (**) | 0.1887 (ns) | <0.0001 (****) | 0.3763 (ns) | 0.0118 (*) | 0.0017 (**) |
| 64x64 | 0.0092 (**) | 0.2205 (ns) | <0.0001 (****) | 0.3549 (ns) | 0.0147 (*) | 0.0019 (**) |

**Table SI.1.** Comparative analysis of the sensitivity of HT29-MTX/HT29-Luciferase HET-tumors exhibiting high MUC1 and EPCAM expression (MUC1[hi]/EPCAM[hi]) at varying grid sizes. The table represents the adjusted P values derived from comparisons between different Antibody-Protein L-MPs. NS (not significant) and asterisks indicate varying levels of significance from * $p < 0.05$, ** $p < 0.01$, *** $p < 0.001$, to **** $p < 0.0001$. Grid sizes range from 8x8 to 64x64.
